# A phosphorylation-controlled switch confers cell cycle-dependent protein relocalization

**DOI:** 10.1101/2024.06.05.597552

**Authors:** Xiaofu Cao, Shiying Huang, Mateusz M. Wagner, Yuan-Ting Cho, Din-Chi Chiu, Krista M. Wartchow, Artur Lazarian, Laura Beth McIntire, Marcus B. Smolka, Jeremy M. Baskin

## Abstract

Tools for acute manipulation of protein localization enable elucidation of spatiotemporally defined functions, but their reliance on exogenous triggers can interfere with cell physiology. This limitation is particularly apparent for studying mitosis, whose highly choreographed events are sensitive to perturbations. Here we exploit the serendipitous discovery of a phosphorylation-controlled, cell cycle-dependent localization change of the adaptor protein PLEKHA5 to develop a system for mitosis-specific protein recruitment to the plasma membrane that requires no exogenous stimulus. Mitosis-enabled Anchor-away/Recruiter System (MARS) comprises an engineered, 15-kDa module derived from PLEKHA5 capable of recruiting functional protein cargoes to the plasma membrane during mitosis, either through direct fusion or via GFP–GFP nanobody interaction. Applications of MARS include both knock sideways to rapidly extract proteins from their native localizations during mitosis and conditional recruitment of lipid-metabolizing enzymes for mitosis-selective editing of plasma membrane lipid content, without the need for exogenous triggers or perturbative synchronization methods.

## INTRODUTION

Extracellular cues are converted into intracellular signaling cascades via regulatory mechanisms that frequently involve conditional recruitment of key enzymes to the plasma membrane (PM) to access substrates, activators, and/or inhibitors^1,2^. Synthetic biology approaches can short-circuit these natural systems, bypassing cell-surface receptors by enabling spatiotemporal control over the localizations and activities of these signaling enzymes. Most commonly, these tools involve conditional protein dimerization following an exogenous trigger such as a small molecule (chemogenetics) or visible light (optogenetics)^3–6^.

Though these systems are widely applied, their limitations include, for chemogenetics, lack of reversibility^7^ and undesired physiological effects of rapamycin^8^, the most frequently used trigger, or, for optogenetic tools, phototoxicity, loss of microscopy channels for observation, and challenges with multiplexing^4,9^. Critically, these methods all involve application of an exogenous perturbation, adding complexity and motivating searches for new triggers (e.g., heat^10^). An alternate approach for non-perturbative conditional protein activation would involve no exogenous trigger at all, instead harnessing an endogenous stimulus.

A natural system in all dividing eukaryotic cells exhibiting strict temporal control over protein activation and deactivation is the cell cycle, which features regulated changes in protein posttranslational modification and proteolysis caused by the oscillatory activities of kinases, phosphatases, and ubiquitin ligases^11–13^. Indeed, the oscillating patterns of enzymatic activities and protein levels have been exploited as endogenous triggers that control levels of different fluorescent proteins to enable visualization of cell cycle stage^14–16^. However, this approach only allows for observation, not manipulation, of cellular processes.

Here, we report the design, optimization, and application of a Mitosis-enabled Anchor-away/Recruiter System (MARS) that enables conditional localization of target proteins to the PM exclusively during mitosis. MARS comprises an engineered 129 amino acid (15 kDa) module from pleckstrin homology domain-containing A5 (PLEKHA5) that localizes to the cytosol during interphase and the PM during mitosis. We describe the discovery of the cell cycle-dependent localization of PLEKHA5 and the engineering of the minimal PLEKHA5-derived MARS module. We then elucidate the mechanism underlying the mitosis-dependent PM localization of PLEKHA5, which involves a single-residue phosphorylation/dephosphorylation switch. Finally, we apply MARS to facilitate recruitment of various protein cargoes, including an endogenously tagged protein kinase, a phospholipase, and a lipid kinase. These studies point to roles for excess phosphatidylinositol 3,4,5-trisphosphate (PI(3,4,5)P_3_) at the PM in hindering progression through anaphase. MARS represents the first example of harnessing mitosis as an endogenous stimulus to mediate reversible protein recruitment without exogenous triggers, and we anticipate it will be useful for manipulating mitosis-specific biological activities without the necessity for synchronization.

## RESULTS

### PLEKHA5 localizes to the PM during mitosis via interaction with PI(4,5)P_2_

During interphase, PLEKHA5 localizes to the microtubule cytoskeleton via N-terminal tandem WW domains^17–19^ (Figure 1A). However, we observed that during mitosis, a substantial pool of EGFP-PLEKHA5^FL^ (FL, full-length) localized to the plasma membrane (PM), in addition to the mitotic spindle (Figure 1B). PLEKHA5 contains a putative membrane-binding motif consisting of an amphipathic helix, an arginine- and lysine-rich basic peptide, and a pleckstrin homology (PH) domain (Figure 1A). This motif (H-BP-PH) shares homology with a motif in the PLEKHA5 paralog PLEKHA4, which localizes to the PM via binding to phosphatidylinositol 4,5-bisphosphate (PI(4,5)P_2_)^20^.

**Figure 1.**
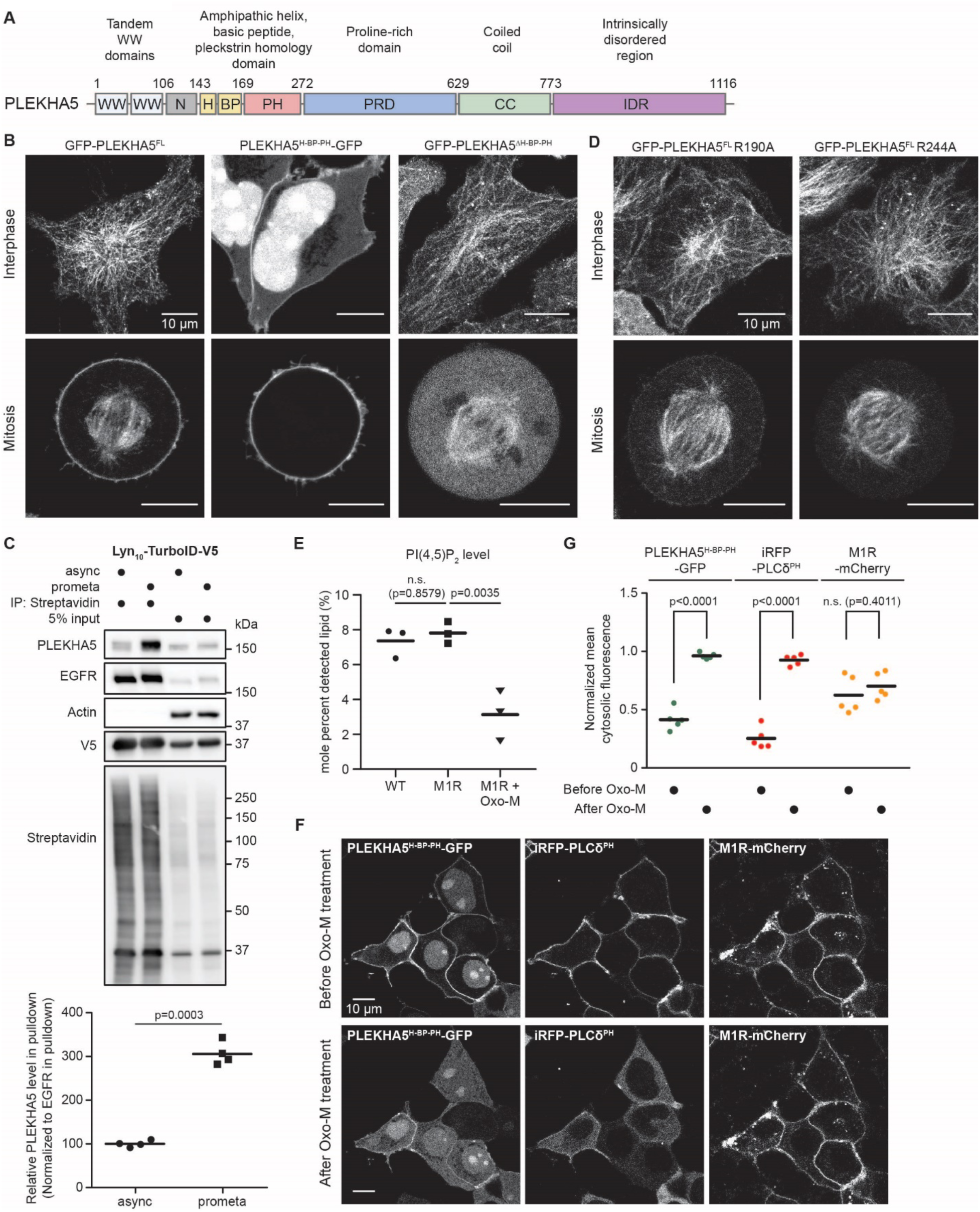
PLEKHA5 exhibits increased PM localization during mitosis via PI(4,5)P_2_ interaction. (A) PLEKHA5 domain map. (B) Live-cell confocal microscopy of HeLa cells transfected with GFP-tagged PLEKHA5 constructs during interphase (top) and mitosis (bottom). (C) A larger portion of endogenous PLEKHA5 was biotinylated by the PM-targeted TurboID construct in isolated prometaphase cells than in the asynchronous populations. Representative western blots are shown at top with quantifications at bottom (n=4 biological replicates, unpaired, two-tailed Student’s t-test). (D) Confocal microscopy of PI(4,5)P_2_ binding mutants (R190A and R244A) of EGFP-PLEKHA5^FL^. (E) Quantification of PI(4,5)P_2_ levels in wild-type (WT) and M1R-mCherry-transfected HeLa cells with or without treatment with the M1R agonist oxotremorine-M (Oxo-M, 30 s) using High Performance Ion Chromatography. (n = 3 biological replicates, one-way ANOVA with Tukey post hoc multiple comparisons test). (F-G) Acute PI(4,5)P_2_ depletion assay in cells co-transfected with M1R-mCherry, the PI(4,5)P_2_ reporter iRFP-PLC8^PH^, and PLEKHA5^H-BP-PH^-GFP reveals the PI(4,5)P_2_-binding capacity of PLEKHA5^H-BP-PH^-GFP. Shown are representative micrographs (F) and quantification (G) of normalized mean fluorescence intensity in the cytosol before and after treatment with Oxo-M (n=5 cells from two independent experiments, unpaired, two-tailed Student’s t-test).

Indeed, GFP-tagged PLEKHA5^H-BP-PH^ exhibited constitutive PM localization during interphase and mitosis with increased PM localization during M phase, similar to FL protein, whereas a PLEKHA5 construct lacking this motif, GFP-PLEKHA5^ΔH-BP-PH^, showed no PM interaction (Figure 1B), demonstrating that the H-BP-PH motif is sufficient and necessary to recruit PLEKHA5 to the PM. Further, the longer N-terminal fragment WW-H-BP-PH exhibited similar localizations as the full-length protein, with little PM association during interphase but increased PM localization during mitosis, establishing the minimal motifs required for the dual localizations of PLEKHA5 during the cell cycle (Extended Data Figure 1A). Extended time-lapse imaging of asynchronous HeLa cells expressing PLEKHA5^WW-H-BP-PH^-GFP and H2B-mCherry (Extended Data Figure 1B and Supplementary Video 1) revealed the first appearance of PM localization at the stage of nuclear envelope breakdown (NEBD), with PM fluorescence peaking in metaphase and then decreasing starting in anaphase and terminating upon completion of cytokinesis, accompanying chromosome decondensation.

To assess the PM localization of endogenous PLEKHA5 during mitosis, we expressed a PM-targeted Lyn_10_-TurboID-V5^21^ fusion to use proximity biotinylation to tag all proteins at or near the PM, followed by streptavidin agarose enrichment and western blot analysis to probe for endogenous PLEKHA5 in the enriched PM-proximal proteome (Extended Data Figure 1C-E).

Indeed, levels of PLEKHA5 biotinylation were higher in prometaphase-synchronized cells relative to asynchronous cells, indicating that endogenous PLEKHA5 also exhibits cell cycle-dependent changes to its PM localization (Figure 1C).

We then tested whether PI(4,5)P_2_ mediates PM binding of the PLEKHA5 H-BP-PH domains, as it does for the paralog PLEKHA4^20^. Accordingly, we found that mutation of two key arginine residues within the PH domain (R190A and R244A), analogous to those required for PI(4,5)P_2_ binding of PLEKHA4 (Extended Data Figure 1F), decreased the PM association of GFP-PLEKHA5, PLEKHA5^WW-H-BP-PH^-GFP, and PLEKHA5^H-BP-PH^-GFP (Figure 1D and Extended Data Figure 1G). Further, acute depletion of PI(4,5)P_2_ by transient pharmacological activation of GPCR/Gq-coupled phospholipase Cs (Figure 1E) led to loss of PM localization of PLEKHA5^H-BP-PH^-GFP (Figure 1F-G), indicating the requirement of PI(4,5)P_2_ for the PM targeting of PLEKHA5.

### Engineering of a mitosis-specific PM-binding motif based on PLEKHA5

Having characterized the dynamic PM localization of PLEKHA5, we sought to engineer a mitosis-specific anchor-away/recruiter system (MARS) based on the H-BP-PH modules. We first had to eliminate any PM localization during interphase while retaining PM association during mitosis. Previously, mutation of four basic residues in the PLEKHA4 BP motif decreased PM localization in cells and PI(4,5)P_2_ binding in vitro^20^. As these residues were conserved in PLEKHA5 (Figure 2A and Extended Data Figure 2A), we analyzed the localizations of several single and double mutants of these residues within PLEKHA5^H-BP-PH^. Only K163A/R164A retained mitotic PM localization while exhibiting nucleocytoplasmic localization in interphase (Figure 2B and Extended Data Figure 2B). Addition of an internal or C-terminal nuclear export signal (NES) eliminated the nuclear localization during interphase (Figure 2C).

**Figure 2.**
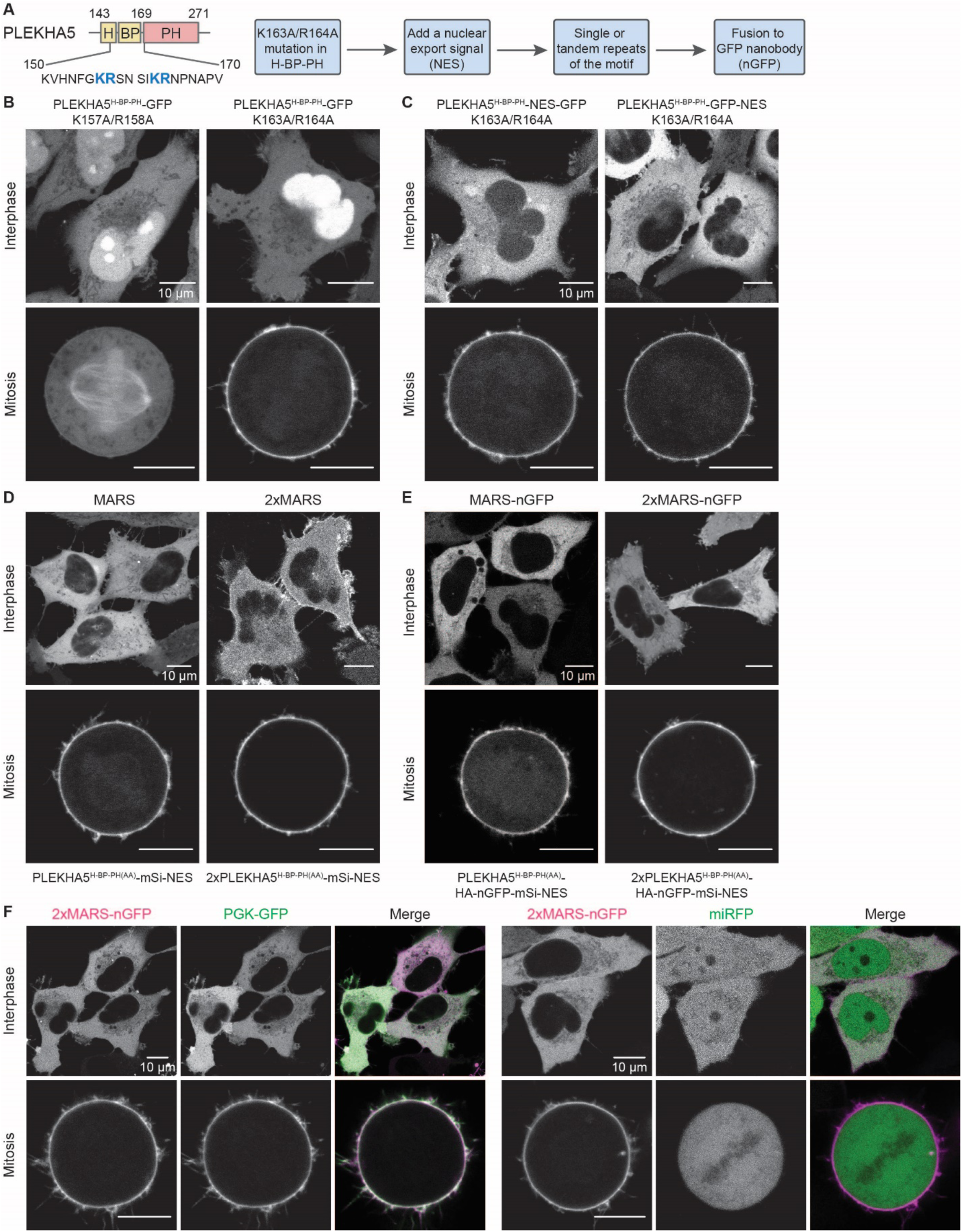
Design and engineering of Mitosis-enabled Anchor-away/Recruiter System (MARS), based on the H-BP-PH motifs of PLEKHA5. (A) Domain map of the PLEKHA5 H-BP-PH motifs with K157, R158, K163, and R164 highlighted in blue, and flow chart of protein engineering workflow to generate MARS. (B) Live-cell imaging of HeLa cells transfected with different mutants of PLEKHA5. (C) Localization of PLEKHA5^H-BP-PH^-GFP with addition of a nuclear export signal (NES) at two different positions. (D-E) Live-cell confocal microscopy to examine subcellular localization of the single and tandem MARS constructs without (D) or with (E) fusion to a GFP nanobody (nGFP). (F) 2xMARS-nGFP is capable of recruiting GFP (left panel) but not miRFP (right panel) to the PM during mitosis.

The module can be fused directly to protein cargoes or to a GFP nanobody (nGFP)^22^ for the recruitment of GFP-tagged proteins. Because of the latter option, we replaced the GFP visualization tag with the red fluorescent protein mScarlet-i (mSi) in the optimized constructs, termed MARS (PLEKHA5^H-BP-PH(AA)^-mSi-NES) and MARS-nGFP (PLEKHA5^H-BP-PH(AA)^-HA-nGFP-mSi-NES). Finally, to enhance avidity for the PM and allow for potential alterations in affinity conferred by cargo fusion/binding, we also constructed tandem repeats termed 2xMARS and 2xMARS-nGFP. Imaging analysis established the expected cytosolic-to-PM shift in localization between interphase and mitosis (Figures 2D-E, Extended Data Figures 2C-D, and Supplementary Videos 2-3). Finally, we demonstrated the functionality of 2xMARS-nGFP, which recruited soluble GFP but not monomeric infrared fluorescent protein (miRFP) to the PM during mitosis (Figure 2F).

We next used quantitative Western blot and confocal microscopy analysis to determine the stoichiometry of GFP to 2xMARS-nGFP required for PM recruitment of GFP (Extended Data Figure 3A). These studies revealed that minimal PM recruitment of GFP was observed when the GFP-to-2xMARS-nGFP molar ratio exceeded 20, whereas partial recruitment occurred at a ratio of ∼14 and complete recruitment required ratios <1.8 (Extended Data Figure 3B-E). These results are aligned with the reported 1:1 stoichiometry of GFP:nGFP complex^22^ and indicate that fusion of 2xMARS to nGFP did not appear to alter its interaction with GFP.

To evaluate any potential disruption to cell cycle progression caused by ectopic expression of MARS or 2xMARS-nGFP, we quantified the proliferation rate of HeLa cells stably expressing MARS and 2xMARS-nGFP compared to untransduced control wild-type HeLa cells using an IncuCyte Live Cell Analysis System. All three cell lines demonstrated similar doubling times during exponential growth phase (Extended Data Figure 3F-G and Supplementary Videos 4-6), indicating that overexpression of these constructs did not perturb rates of cell proliferation.

### The PM localization of PLEKHA5 is regulated by the phosphorylation status of S161

We next investigated the underlying molecular mechanism governing the dynamic PM localization of PLEKHA5. Because PI(4,5)P_2_ levels do not increase during mitosis in HeLa cells^23^, we hypothesized that posttranslational modifications to PLEKHA5 might regulate its localization, in particular phosphorylation^24,25^. Therefore, we generated phosphodeficient and phosphomimetic mutations of nine residues that were adjacent to the PH domain and reported to be phosphorylated in PhosphoSitePlus^26^, within both GFP-PLEKHA5^FL^ and PLEKHA5^WW-H-BP-PH^-GFP. Mutations at eight positions caused no change in localization by confocal microscopy (Extended Data Figure 4). Strikingly, however, mutation of S161 strongly affected the subcellular localization. The S161A phosphodeficient mutants of both GFP-PLEKHA5^FL^ and PLEKHA5^WW-H-BP-PH^-GFP displayed increased PM association during interphase, whereas the S161D phosphomimetic mutants exhibited no PM localization during mitosis (Extended Data Figures 5A-D). We also used TurboID-based proximity labeling of the PM-associated proteome to confirm the changes in localization observed by microscopy (Extended Data Figures 5E-F). The S161 mutations were also incorporated into the PLEKHA5^H-BP-PH^-GFP, and imaging analyses demonstrated the expected changes in PM localization (Figure 3B).

**Figure 3.**
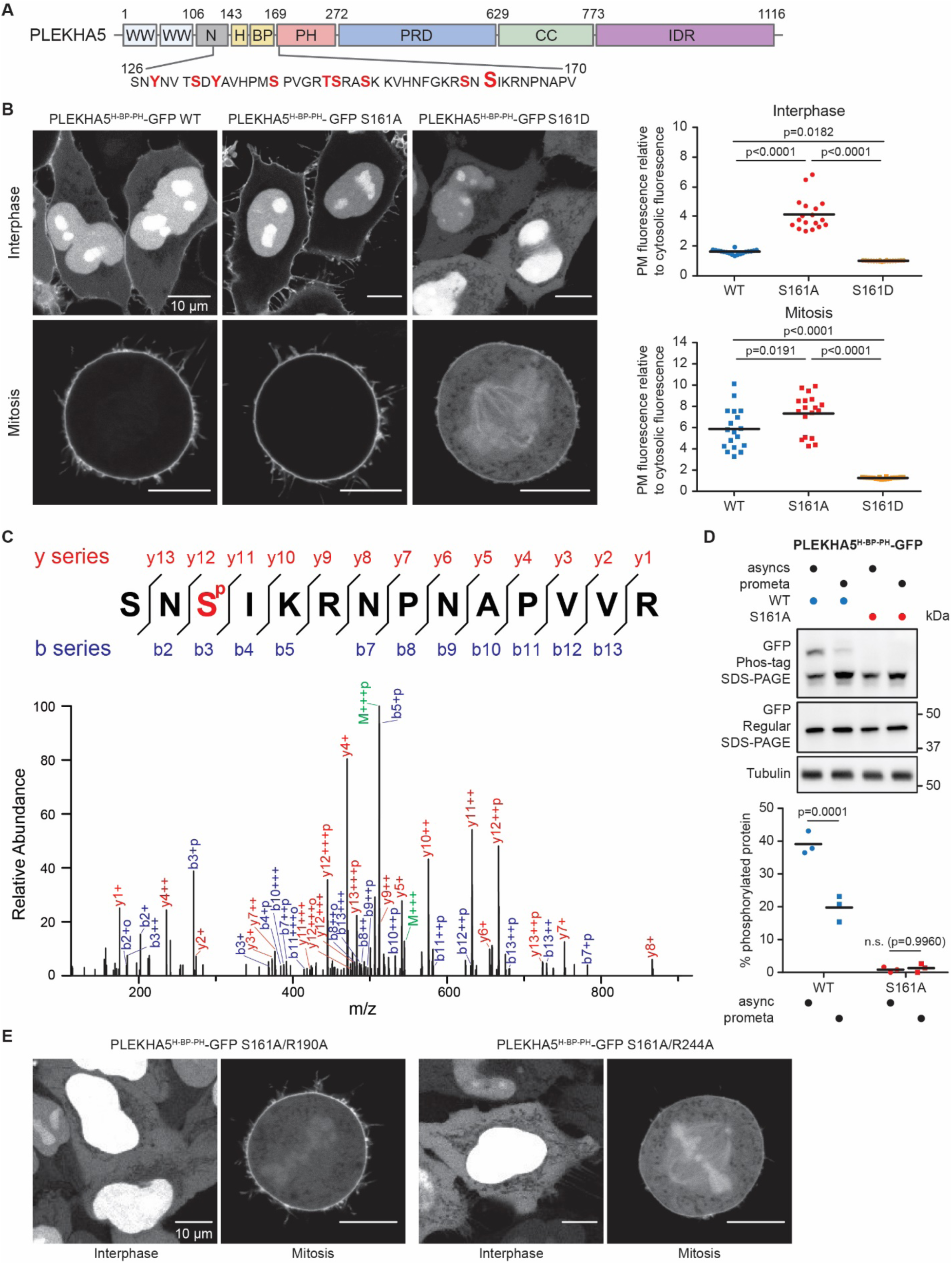
The PM localization of PLEKHA5 is regulated by S161 phosphorylation. (A) Primary amino acid sequence of the region of PLEKHA5 immediately upstream of the PH domain. Residues highlighted in red were selected for phospho-mutagenesis analysis. (B) The S161A mutation increased the PM localization of PLEKHA5^H-BP-PH^-GFP, whereas the S161D mutation diminished the PM association of this construct. Shown at left are representative micrographs and at right the quantification of PM to cytosolic fluorescence ratios in HeLa cells transfected with each construct in interphase and mitosis (n=18 cells from three independent experiments, one-way ANOVA with Tukey post hoc multiple comparisons test). (C) MS^2^ spectrum of phosphoproteomics analysis of GFP immunoprecipitates from asynchronous HeLa cells transfected with PLEKHA5^H-BP-PH^-GFP. Labeled peaks show fragmented phosphate-containing ions. “o” following the charge descriptor refers to a neutral water loss; “p” refers to water and phosphate loss. “M” marks precursor ions. The sequence of the detected mis-cleaved peptide from tryptic digest with corresponding b and y series ions is shown at top. The S161 site is highlighted in red. (D) Phos-tag and regular SDS-PAGE analysis of lysates harvested from asynchronous or prometaphase-synchronized HeLa cells transfected with WT or S161A forms of PLEKHA5^H-BP-PH^-GFP. Shown are representative western blots (top) and quantification of phosphorylated proteins (bottom) (n=3 biological replicates, one-way ANOVA with Tukey post hoc multiple comparisons test). (E) Live-cell imaging of HeLa cells transiently expressing combinational mutant of S161A with either R190A or R244A for PLEKHA5^H-BP-PH^-GFP illustrating the requirement for PI(4,5)P_2_ binding for PM association independent of S161 phosphorylation status.

Because these data indicate that the S161A mutation enhances PM binding and the S161D mutation eliminates such localization, these results suggest that phosphorylation at S161, mimicked by S161D, might prevent the PM localization of PLEKHA5 during interphase and that dephosphorylation during mitosis, mimicked by S161A, might enable PM binding.

First, to confirm the phosphorylation of PLEKHA5 at S161, we performed phosphoproteomics analysis of enriched PLEKHA5^H-BP-PH^-GFP and detected a miscleaved tryptic phosphopeptide with MS^2^ analysis confirming phosphorylation at S161 (Figure 3C). We also exploited Phos-tag SDS-PAGE^27^, wherein phosphorylated forms of a given protein migrate more slowly than the non-phosphorylated forms, to interrogate the mechanisms underlying S161 phosphorylation. Analysis of lysates from asynchronous or prometaphase-synchronized cells expressing either wild-type (WT) or S161A forms of PLEKHA5^H-BP-PH^-GFP revealed a loss of a slower-migrating band in the S161A samples that likely correspond to PLEKHA5^H-BP-PH^-GFP phosphorylated at S161 (Figure 3D). The intensity of this band in WT PLEKHA5^H-BP-PH^-GFP was significantly lower in prometaphase-synchronized compared to asynchronous cells, consistent with the phosphomutant imaging results suggesting that a larger portion of PLEKHA5^H-BP-PH^-GFP was dephosphorylated during mitosis.

To further understand how S161 phosphorylation modulates the PM association of PLEKHA5 and MARS, we analyzed the localizations of the PI(4,5)P_2_ binding deficient mutants, R190A and R244A, in the S161A background. The S161A mutation did not restore the PM localizations of GFP-PLEKHA5^FL^, PLEKHA5^WW-H-BP-PH^-GFP, or the PLEKHA5^H-BP-PH^-GFP constructs harboring either R190A or R244A (Figure 3E and Extended Data Figure 6A-B), implying that the S161 phosphorylation switch on the BP requires an intact PI(4,5)P_2_-binding PH domain to regulate PM association.

Interestingly, the PLEKHA5 paralog PLEKHA4 exhibits plasma membrane localization in both interphase^20^ and mitosis (Extended Data Figure 6C), and a sequence alignment of the BP motifs from these proteins revealed that human PLEKHA4 contains an alanine at position 46, which corresponds to S161 in human PLEKHA5 (Extended Data Figure 6D). Examination of the localization of A46S and A46D mutants of GFP-PLEKHA4 revealed that, though the A46S mutant exhibited a similar PM localization to the WT protein, the A46D mutation abolished the PM localization during both interphase and mitosis (Extended Data Figure 6E). These results suggest the importance of this residue in regulating the PM binding of protein members across the PLEKHA sub-family.

### PKC and CK1 kinases mediate PLEKHA5 S161 phosphorylation

Next, we sought to determine the kinases responsible for PLEKHA5 S161 phosphorylation. According to a kinase/phosphosite prediction tool^28^, several protein kinase C (PKC) isoforms were among the top predicted kinases for recognizing the linear amino acid sequence around S161, prompting us to examine the PKC family (Figure 4A). We first treated cells expressing either the WT or S161A forms of PLEKHA5^H-BP-PH^-GFP with one of two PKC agonists, phorbol 12-myristate 13-acetate (PMA)^29^ or Bryostatin 1^30^, or a DMSO control, with or without pretreatment with a pan-PKC inhibitor cocktail (PKCi: Gö 6983 and Sotrastaurin/AEB071)^31–33^. Subsequent Phos-tag SDS-PAGE analysis revealed that treatment with either PKC agonist significantly increased the intensity of the slower-migrating, phospho-S161 band, and that this effect was eliminated in the presence of PKC inhibition (Figure 4B). Furthermore, live-cell, time-lapse imaging of both PLEKHA5^H-BP-PH^-GFP and PLEKHA5^WW-H-BP-PH^-GFP upon PKC activation revealed that addition of PMA or Bryostatin 1 induced a loss of PM localization within 25 minutes, an effect that was abolished upon pre-treatment of PKC inhibitors or by using S161A mutants (Figures 4C-D and Extended Data Figures 7A-B).

**Figure 4.**
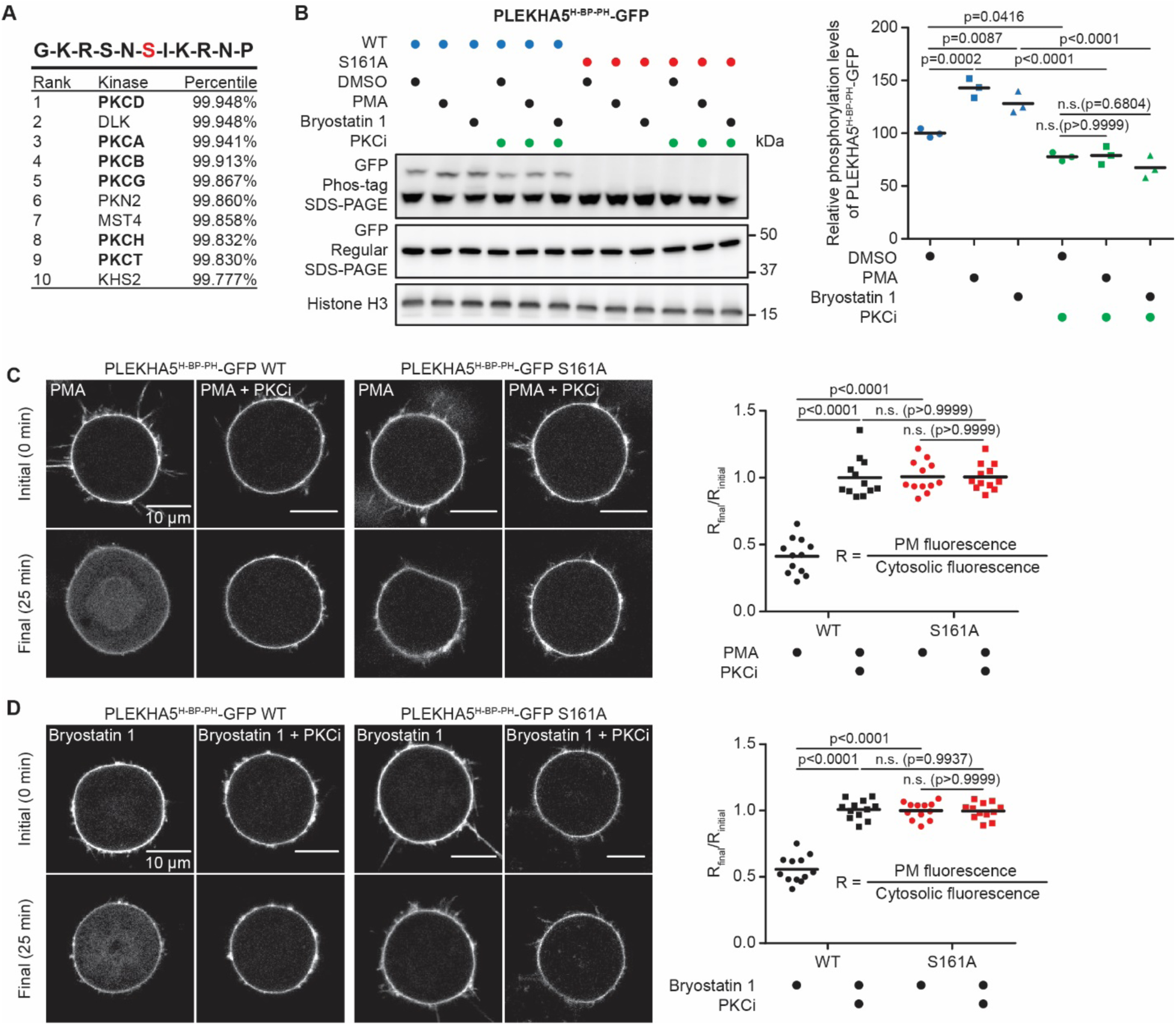
S161 phosphorylation and PLEKHA5 PM localization are modulated by protein kinase C (PKC) activity. (A) Top 10 kinase candidates for phosphorylating S161. PKC isoforms are bolded. (B) S161 phosphorylation increased upon stimulation of PKC activity using 100 nM PMA or 200 nM Bryostatin 1, and the increase was attenuated with pre-treatment of PKC inhibitors (PKCi, 100 nM Gö 6983 and 100 nM Sotrastaurin/AEB071). Representative western blots (left) and quantification (right) from Phos-tag and regular SDS-PAGE analysis probing PLEKHA5^H-BP-PH^-GFP are shown (n=3 biological replicates, one-way ANOVA with Sidak post hoc multiple comparisons test). (C-D) PM localization of PLEKHA5^H-BP-PH^ was decreased upon activation of PKC by 100 nM PMA (C) or 200 nM Bryostatin 1 (D). Representative confocal micrographs before and after agonist treatment for each condition are shown at left, with quantification of the ratios of PM to cytosolic fluorescence before and after treatment are shown at right (n=12 cells from three independent experiments, one-way ANOVA with Sidak post hoc multiple comparisons test).

Though phosphodeficient/mimetic mutants of the nearby S159 did not affect PLEKHA5 PM localization, this residue is reported to be phosphorylated^26^, and a different set of kinases were predicted to mediated S161 phosphorylation in the presence of phospho-S159, notably several casein kinase 1 (CK1) isoforms^28^ (Extended Data Figure 7C). Indeed, Phostag SDS-PAGE analysis revealed that CK1 inhibition by PF670462^34–36^ or D4476^37^ significantly decreased the level of S161 phosphorylation of PLEKHA5^H-BP-PH^-GFP, and combinational treatment with both CK1 and PKC inhibitors eliminated the majority of S161 phosphorylation (Extended Data Figure 7D). Collectively, these data point to the PKC and CK1 kinase families as important mediators of PLEKHA5 and MARS S161 phosphorylation during interphase and imply that such phosphorylation activity toward PLEKHA5 S161 is reduced during mitosis.

Next, we examined whether the PPP family of phosphoprotein phosphatases dephosphorylate PLEKHA5 phospho-S161, because members of this serine/threonine phosphatase family carry out numerous dephosphorylation events related to mitosis^38^. Pan inhibition of PPP phosphatases using okadaic acid^39,40^ induced elevation of phospho-S161 levels, but treatment of PP1- or PP2A/4-selective inhibitors (Tautomycetin^39^ and Fostriecin^41^, respectively) caused minimal effects on S161 phosphorylation, indicating the involvement of multiple phosphatases in this dephosphorylation event (Extended Data Figure 7E).

Finally, we independently validated the effects of these pharmacological treatments on S161 phosphorylation by instead using siRNA knockdown of the major isoforms of PKC and CK1 and the catalytic subunits of PP1 and PP2A expressed in these cells (Extended Data Figure 7F). Phostag SDS-PAGE revealed that knockdown of CK1 or PKC caused reduction in S161 phosphorylation of PLEKHA5^H-BP-PH^-GFP, whereas depletion of PP1 and PP2A led to an increase in S161 phosphorylation (Extended Data Figure 7G).

### MARS can recruit multiple functional enzymatic cargoes to the PM during mitosis

We next evaluated the general ability of MARS to mediate mitosis-conditional recruitment of functional enzyme cargoes to the PM (Figure 5A). First, we selected polo-like kinase 1 (PLK1), a critical regulator of mitosis with multiple important non-PM localizations during both interphase and mitosis. We expressed 2xMARS-nGFP, chosen over MARS-nGFP due to its low cytoplasmic background during mitosis (Figure 2E), in cells expressing endogenously tagged PLK1-GFP. Control cells transfected with nGFP-free MARS exhibited expected PLK1-GFP localization at centromeres, midbody, mitotic kinetochores, and spindle poles^42^ (Figure 5B). Expression of 2xMARS-nGFP, however, induced PM recruitment of a substantial pool of PLK1-GFP exclusively during mitosis and not during interphase (Figure 5B). However, during interphase, a pool of PLK1-GFP was recruited from the nucleus to the cytosol due to its interaction with 2xMARS-nGFP (Figure 5B). This observation points to a limitation of the system, namely that colocalization of GFP cargoes with 2xMARS-nGFP persists throughout the cell cycle. Thus, MARS systems that bear an NES are most compatible with cytosolic proteins, though a version of MARS without an NES (e.g., K163A/R164A mutants of PLEKHA5^H-BP-PH^) exhibits a more prominent nuclear localization (Figure 2B) and thus may be more useful for proteins with nuclear or nucleocytoplasmic localizations.

**Figure 5.**
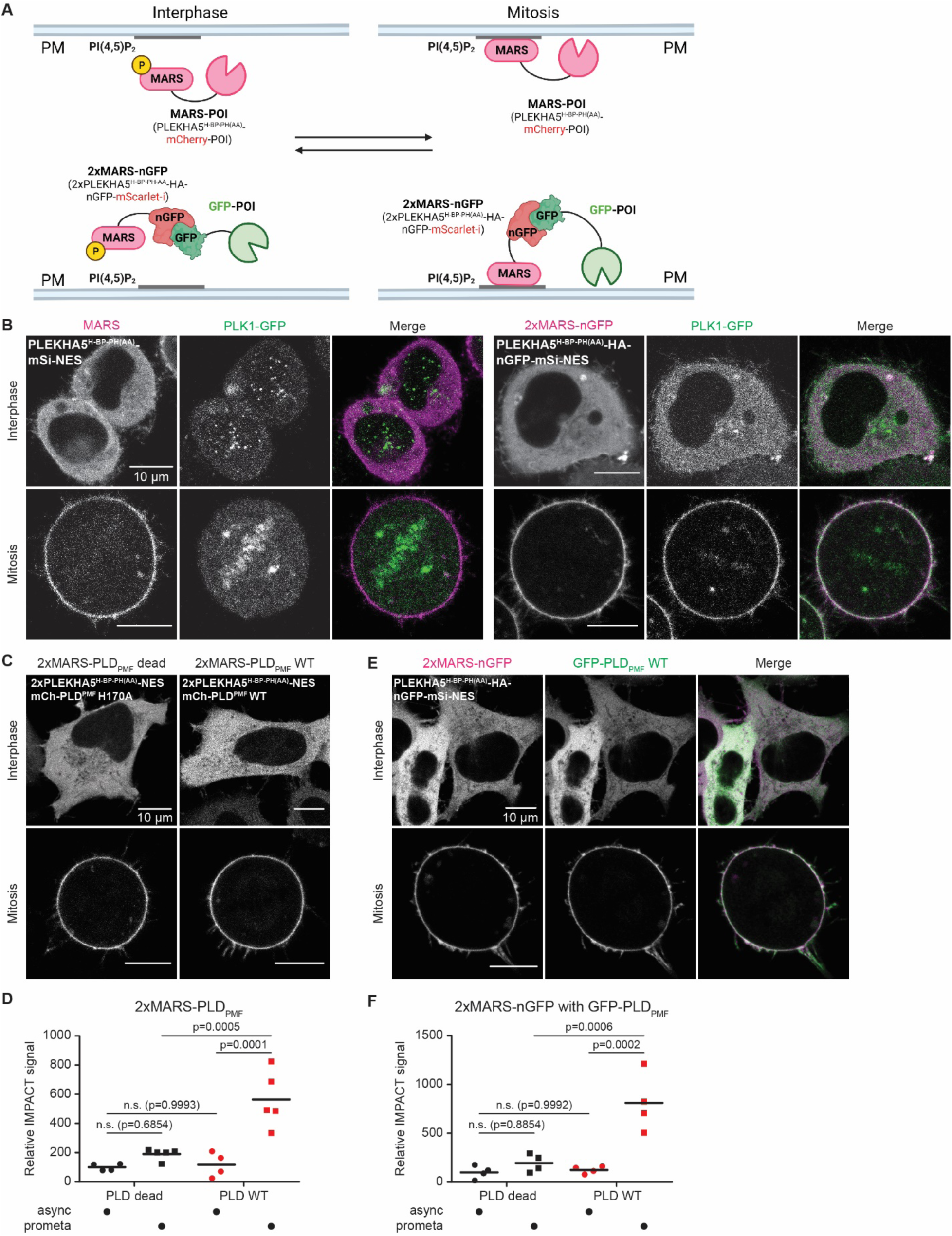
MARS recruits polo-like kinase 1 (PLK1) and phospholipase D (PLD) to the PM specifically during mitosis. (A) Cartoon illustration of the working principle of MARS for either direct fusion (top) or nGFP-based recruitment (bottom) of proteins of interest (POI) to the PM during mitosis. Cartoon is generated using BioRender. (B) 2xMARS-nGFP (magenta, transient transfection) anchored PLK1-GFP (green, endogenously tagged) at the PM during mitosis in HeLa cells. (C-D) Tandem MARS fusion to PLD from *Streptomyces sp.* PMF (PLD_PMF_) enabled mitosis-induced PM recruitment of functional PLD_PMF_. (C) Live-cell confocal microscopy of HeLa cells transfected with catalytically dead (H170A) or WT 2xMARS-PLD_PMF_ in interphase and mitosis. (D) Measurements of relative activity of 2xMARS-PLD_PMF_ variants in asynchronous or isolated prometaphase cells measured by IMPACT labeling followed by flow cytometry demonstrate elevated activity of WT but not catalytically dead PLD concomitant with its selective recruitment to the PM during mitosis (n=4 biological replicates for asynchronous cell, and n=5 biological replicates for prometaphase cells, one-way ANOVA with Sidak post hoc multiple comparisons test). (E-F) 2xMARS-nGFP recruited active GFP-PLD_PMF_ to the PM during mitosis. (E) Representative confocal micrographs of HeLa cells transfected with 2xMARS-nGFP (magenta) and WT GFP-PLD_PMF_ (green). (F) Measurements of relative activity of GFP-PLD_PMF_ variants in asynchronous or isolated prometaphase cells measured by IMPACT labeling followed by flow cytometry demonstrate elevated activity of WT but not catalytically dead PLD concomitant with its selective recruitment to the PM during mitosis (n=4 biological replicates, one-way ANOVA with Sidak post hoc multiple comparisons test).

We next tested a phospholipase D from *Streptomyces sp.* PMF (PLD_PMF_), which converts the abundant phospholipid phosphatidylcholine (PC) to the signaling lipid phosphatidic acid (PA) and which we have harnessed for use as an optogenetic membrane editor for controlled phospholipid modification in mammalian cells^43–45^. Direct fusion of 2xMARS to PLD_PMF_ (with mCherry in place of mSi) led to complete PM recruitment during mitosis (Figure 5C), whereas fusion of a single MARS motif led to only partial PM recruitment (Extended Data Figure 8A). We then assessed whether such PLD recruitment induced membrane editing of PC to PA in mitosis by measuring the enzymatic activity of 2xMARS-PLD_PMF_ in asynchronous or mitotic cells using our previously developed chemoenzymatic method, IMPACT, coupled to flow cytometry^29,44^.

Indeed, compared to asynchronous cells, 2xMARS-PLD_PMF_ exhibited significantly higher IMPACT fluorescence in prometaphase cells, when it was recruited to the PM, and a catalytically dead form (bearing the H170A mutation in PLD_PMF_) exhibited low IMPACT signal at all cell cycle stages, demonstrating the specificity of the IMPACT assay to MARS-associated PLD activity (Figure 5D). These results show that direct fusion of 2xMARS to PLD_PMF_ does not impair its catalytic function and permits mitosis-specific PM recruitment. Although quantification of IMPACT fluorescence by flow cytometry reports on whole-cell PLD activity, the increase in IMPACT seen only in mitotic cells expressing catalytically active 2xMARS-PLD_PMF_ indicates its utility as a reporter of PM-localized PLD activity in this context. Alternatively, the use of 2xMARS-nGFP to recruit exogenously expressed GFP-PLD_PMF_ to the PM during mitosis was also successful (Extended Data Figure 8B and Figure 5E). Again, PLD_PMF_ exhibited mitosis-specific membrane editing activity as assessed by IMPACT (Figure 5F), illustrating effective recruitment of a functional GFP-tagged lipid-modifying enzyme by 2xMARS-nGFP.

A third protein cargo we applied MARS to was Class I phosphoinositide 3-kinase (PI3K), which phosphorylates PI(4,5)P_2_ to form PI(3,4,5)P_3_^46^. Following precedent for optogenetic PI3Ks^47^, we fused MARS to an mCherry-tagged inter-Src homology 2 (iSH2) domain of the PI3K regulatory subunit p85 (MARS-p85^iSH2^), which can heterodimerize the endogenous PI3K catalytic subunit, p110^4^. MARS-p85^iSH2^ expression recruited a functional Class I PI3K to the PM and induced formation of PI(3,4,5)P_3_ at this membrane only during mitosis, visualized using the PI(3,4,5)P_3_ biosensor GFP-Akt^PH^, with MARS-p85^iSH2^-induced PI(3,4,5)P_3_ levels significantly higher than basal PI(3,4,5)P_3_ levels in control cells expressing MARS only (Figures 6A-B). In contrast, the tandem 2xMARS-p85^iSH2^ exhibited undesired PM association during interphase, illustrating the importance of having both the single and tandem constructs as options for MARS-cargo design (Extended Data Figure 8C). Again, the nGFP-based system worked as an alternate method for PM recruitment of PI3K during mitosis. Here, we co-expressed GFP-p85^iSH2^ or GFP only with 2xMARS-nGFP and a PI(3,4,5)P_3_ biosensor, Akt^PH^-miRFP670 and observed expected mitotic cargo recruitment for GFP-p85^iSH2^ and GFP but PI(3,4,5)P_3_ synthesis only upon mitotic GFP-p85^iSH2^ recruitment (Figures 6C-D). These studies demonstrate that both direct fusion and the nGFP-based system can enable conditional PI3K recruitment and PI(3,4,5)P_3_ synthesis at the PM during mitosis.

**Figure 6.**
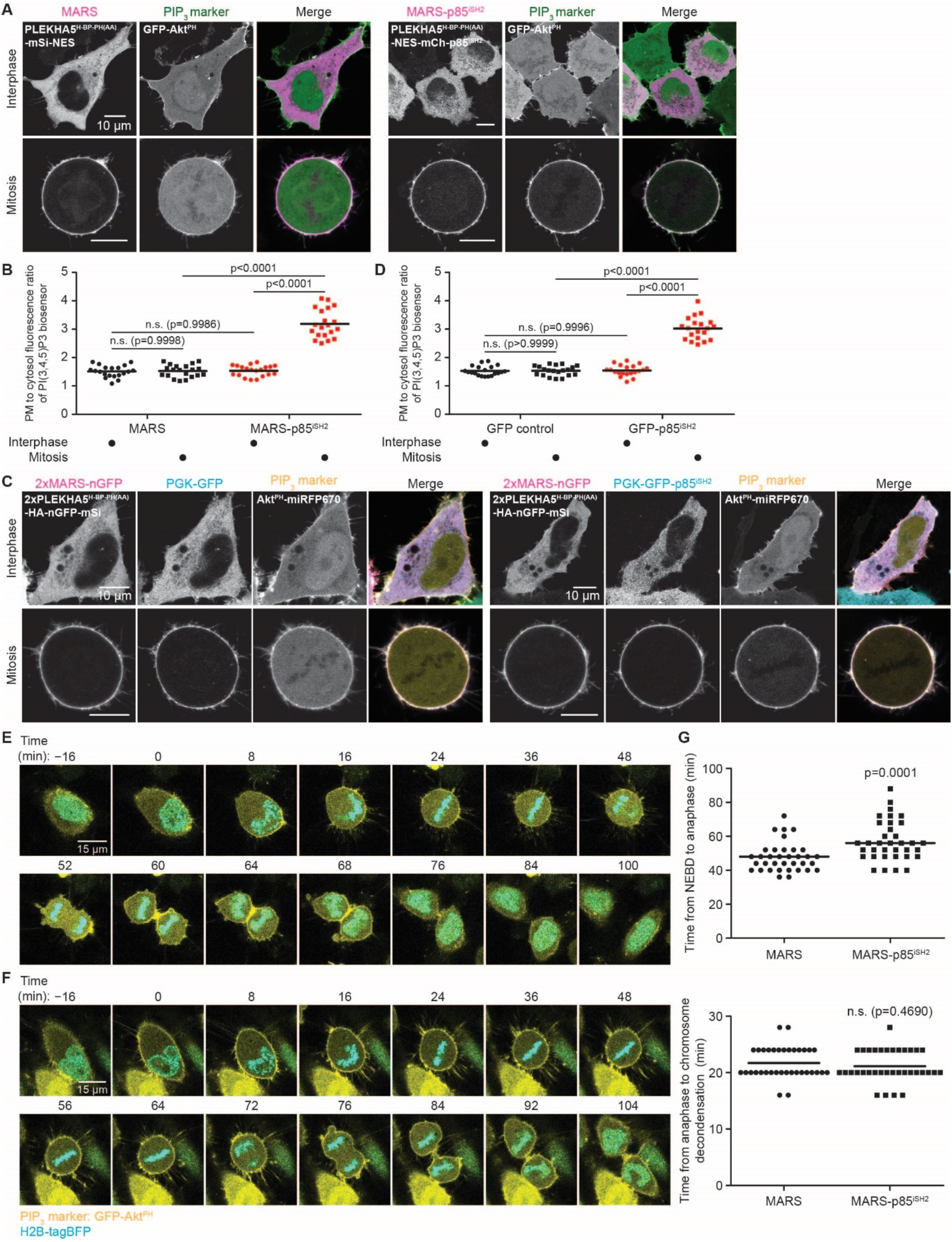
MARS-mediated mitotic PM recruitment of Class I PI3K reveals effects of excessive PI(3,4,5)P_3_ production at the PM during mitosis. (A-B) Single MARS fusion to p85^iSH2^ was sufficient to recruit endogenous PI3K to the PM to generate PI(3,4,5)P_3_ during mitosis. (A) Confocal microscopy of HeLa cells transiently expressing MARS (control, magenta) or MARS-p85^iSH2^ (magenta) along with the PI(3,4,5)P_3_ biosensor GFP-Akt^PH^ (green). (B) Quantification of the PM to cytosolic fluorescence ratios of GFP-Akt^PH^ as a readout for measuring PM PI(3,4,5)P_3_ levels (n=20 cells from three independent experiments, ANOVA with Sidak post hoc multiple comparisons test). (C-D) 2xMARS-nGFP recruited GFP-p85^iSH2^ to the PM during mitosis and produced PI(3,4,5)P_3_. (C) Representative micrographs of HeLa cells co-transfected with 2xMARS-nGFP (magenta), PGK-GFP or PGK-GFP-p85^iSH2^ (cyan), and an miRFP-tagged PI(3,4,5)P_3_ biosensor, Akt^PH^-miRFP670 (yellow). (D) Quantification of the PM to cytosolic fluorescence ratios of Akt^PH^-miRFP670 as the measure for PM PIP_3_ level (n = 20 cells from three independent experiments, ANOVA with Sidak post hoc multiple comparisons test). (E-G) Excess PM PI(3,4,5)P_3_ levels during mitosis caused a delay in mitosis progression through anaphase. (E-F) Representative still images from time-lapse movies in asynchronous HeLa cells stably expressing H2B-tagBFP and transfected with either MARS (E) or MARS-p85^iSH2^ (F), and GFP-Akt^PH^. Images display merged channels of H2B-tagBFP (cyan) and GFP-Akt^PH^ (yellow). Numbers indicate time before/after nuclear envelope breakdown (defined as t=0 min). (G) Quantification of mitosis progression in asynchronous HeLa cells stably expressing H2B-tagBFP and transfected with GFP-Akt^PH^ and either MARS or MARS-p85^iSH2^, with time from NEBD to chromosome separation at anaphase onset (top) and from anaphase onset to chromosome decondensation in cytokinesis (bottom) plotted as scatter plots (n=35 cells from two independent experiments, two-tailed Mann–Whitney U test).

### MARS-p85^iSH2^ reveals an inhibitory effect of a PM PI(3,4,5)P_3_ pool on early mitosis progression

PM-localized phosphoinositides (PI4P, PI(4,5)P_2_, and PI(3,4,5)P_3_) play several key roles throughout cell division, including cell rounding beginning in prophase, spindle orientation during metaphase, and furrow ingression during cytokinesis^48^. These functions have been investigated largely by genetic, pharmacological, and chemogenetic methods for chronically or acutely manipulating PIP levels that are often employed in conjunction with harsh cell synchronization methods that can perturb physiology^49–51^. MARS presents an opportunity to manipulate PM PI(3,4,5)P_3_ levels in bulk cell populations without synchronization, prompting us to investigate whether MARS-p85^iSH2^ might reveal new mitotic roles for PI(3,4,5)P_3_.

We performed live-cell, time-lapse microscopy of HeLa cells expressing H2B-tagBFP to mark chromosomes, GFP-Akt^PH^ to visualize PI(3,4,5)P_3_, and either MARS-p85^iSH2^ or MARS as a negative control, over 14 hours. As expected, MARS and MARS-p85^iSH2^ exhibited PM localization from NEBD until telophase and otherwise cytosolic localization (Extended Data Figure 8D-E, Supplementary Videos 7-8). GFP-Akt^PH^ localization in control cells expressing MARS revealed homeostatic PI(3,4,5)P_3_ pools that did not exhibit significant PM enrichment until telophase and cytokinesis, where fluorescence was observed at the cell periphery and between the two daughter cells, consistent with previous observations (Figure 6E and Supplementary Video 7)^49^. In MARS-p85^iSH2^-expressing cells, GFP-Akt^PH^ displayed elevated PM signal starting from NEBD, confirming mitosis-induced PI(3,4,5)P_3_ production at the PM (Figure 6F and Supplementary Video 8).

Quantification of the duration of the early stages of mitosis, i.e., from NEBD during prophase to chromosomal separation at anaphase onset, revealed a significant increase in the time needed to enter anaphase in cells with excess PI(3,4,5)P_3_ (Figure 6G). However, a similar delay was not observed for mitotic exit, i.e., from anaphase onset to chromosomal decondensation during telophase and cytokinesis (Figure 6G). Therefore, overproduction of PI(3,4,5)P_3_ on the PM during mitosis impairs regular mitosis progression through anaphase, but not afterward, potentially due to improper spindle orientation triggered by the presence of excess PI(3,4,5)P_3_, as first proposed by studies employing chronic perturbations such as PTEN knockdown^49^. These results reveal distinct effects for PM PI(3,4,5)P_3_ at different stages of mitosis and, more broadly, showcase the utility of MARS for conditional modulation of signaling events occurring at the PM during mitosis with high spatiotemporal precision and without synchronization or other exogenous perturbations.

## DISCUSSION

The development of MARS was inspired by the observation that a pool of PLEKHA5 translocates to the PM during mitosis. Mechanistic studies revealed that PI(4,5)P_2_ binding mediates this PM localization but that it is critically regulated by the phosphorylation status of a single residue, S161, whose phosphorylation antagonizes PM recruitment and is decreased during mitosis. Although several kinase families are predicted to phosphorylate S161^28^, we found PKC to contribute substantially to this phosphorylation, raising the question of how S161 dephosphorylation might accompany mitosis onset. Interestingly, two PKC isozymes, PKCα and PKCýII, which were among the most highly predicted kinases for S161 phosphorylation, accumulate in the nucleus during the G2-to-M transition, where they phosphorylate lamin B1 to drive its disassembly and promote NEBD^52–54^. The timing of the nuclear accumulation of these kinases — and therefore their concomitant depletion from the cytoplasm, where PLEKHA5 resides in interphase on the microtubule cytoskeleton — correlates with the onset of PLEKHA5 recruitment to the PM. Therefore, we propose a model wherein reductions in cytoplasmic PKC activity (and potentially that of CK1 and/or additional kinases), coupled to activity of PPP family phosphatases during the G2-to-M transition and in early mitosis, cooperate to reduce S161 phosphorylation and enhance PLEKHA5 PM localization.

Inspired by this dynamic PM localization of PLEKHA5, we engineered a Mitosis-enabled Anchor-away/Recruiter System, MARS, for conditional PM recruitment of protein cargoes during mitosis. MARS is based on a 129 amino acid (15 kDa) PM binding motif from PLEKHA5 but containing additional modifications to ensure exclusive cytosolic localization during interphase and PM localization during mitosis. Versatility is built into MARS in two ways. First, single and tandem (2xMARS) modules were constructed to provide options for low- and high-affinity PM localization, with the tandem modules particularly useful for larger, bulkier cargoes for which fusion to MARS may attenuate the membrane translocation of the fusion construct. Second, MARS can be either fused directly to the protein cargo or a chimeric protein containing MARS fused to a GFP nanobody (MARS-nGFP) that can recruit GFP-tagged proteins. Direct fusion may be more straightforward for modulating signaling via ectopic expression of the protein of interest, whereas the nanobody-based system enables relocalization of endogenously tagged proteins and may be more suited for anchor-away/knock-sideways applications. Collectively, we demonstrated the ability of various MARS constructs to efficiently recruit three different functional cargoes to the PM during mitosis, including an endogenously tagged protein kinase (PLK1) and two lipid-modifying enzymes (PLD and PI3K) capable of editing PM lipid content to produce phosphatidic acid and PI(3,4,5)P_3_ respectively. Our studies in developing mitotic PM recruitable versions of these proteins highlight the importance of both the direct fusion vs. GFP nanobody and single vs. tandem MARS options in construct design.

One potential application of MARS would be for anchor-away of endogenously tagged cytoplasmic proteins with important mitotic functions from their endogenous location(s) to the PM, as we demonstrated for PLK1. Upon mitosis onset, rapid MARS-mediated knock sideways could enable phenotypic and mechanistic investigation downstream of the mitosis-specific loss-of-function, or more accurately loss-of-localization, for these proteins. Removing the NES from MARS would expand its application to nuclear proteins while allowing such cargoes to maintain their proper nuclear localization during interphase. The success of such a strategy would require careful control of expression levels of each component given the stoichiometry of the nGFP–GFP interaction. Additionally, the nGFP–GFP interaction, which would occur constitutively, would have to not be perturbative to the targeted protein during interphase. Finally, to properly interpret results, control experiments would be required to assess the functionality of target proteins fused to MARS or complexed to 2xMARS-nGFP during interphase vs. mitosis using combinations of knockout/knockdown and/or overexpression. Therefore, MARS may represent an approach for functional depletion of enzymes to complement genetic knockdown, pharmacological inhibition, and other stimulus-triggered anchor-away strategies, with the advantage that its onset is highly temporally controlled, and it does not require perturbative synchronization methods for implementation in bulk populations.

A second type of application for MARS is to spatiotemporally modify PM lipid metabolism and signaling pathways during mitosis, e.g., by acute production of two anionic signaling lipids, PA and PI(3,4,5)P_3_. Notably, our studies using MARS to specifically elevate PI(3,4,5)P_3_ levels at the PM during mitosis revealed a selective effect of excess PI(3,4,5)P_3_ in delaying mitotic progression up to the spindle checkpoint, but not afterward. Importantly, these cells were otherwise unperturbed, highlighting the ability of MARS to mediate acute manipulation of lipid metabolism during mitosis with a high degree of spatiotemporal control and without synchronization or other external stimuli. We envision application of MARS for spatiotemporally controlled editing of additional lipids or manipulation of signaling pathways at the PM to investigate mitosis-specific functions. Further, beyond editing of the lipid composition of membranes during mitosis, MARS could also be used for targeted manipulations to protein abundance or localization. For example, PM recruitment of a ubiquitin ligase or other posttranslational modification enzymes could facilitate targeted protein degradation, phosphorylation, or glycosylation of PM pools of proteins of interest^55–60^. Alternatively, PM recruitment of TEV protease during mitosis could be used to enable release of a protein of interest (e.g., a transcription factor) tethered to the PM via a linker containing a TEV substrate sequence to affect downstream signaling and/or transcription, potentially integrating with existing systems that leverage protease cleavage for various applications^61–65^.

In sum, we identify and characterize a phosphorylation/dephosphorylation switch that governs the PM-cytoplasmic cycling of a multidomain adaptor, PLEKHA5, and we then engineer a minimal motif from this protein into a module termed MARS that facilitates selective recruitment of protein cargoes to the PM exclusively during mitosis in otherwise unperturbed cells. MARS represents a versatile and tunable platform for mediating protein anchor-away to rapidly extract proteins from their native localizations to the PM during mitosis, and it can also facilitate conditional recruitment of lipid-metabolizing enzymes to the PM for mitosis-selective membrane editing to precisely manipulate lipid composition and signaling. Given that many proteins exhibit dynamic localization changes that accompany natural oscillations in their posttranslational modification and/or binding interactions with other biological molecules, the self-contained nature of MARS, whose translocation exploits endogenous signaling events, will hopefully inspire the development of additional tools for conditional protein activation as a function of changes to cellular physiological states without need for exogenous triggers.

## MATERIALS AND METHODS

### Antibodies

Mouse anti-PLEKHA5, Santa Cruz Biotechnology (sc-390311, RRID: N/A); rabbit anti-EGFR, Rockland (100-401-149, RRID: AB_2293276); mouse anti-actin, MP Bio (08691001, RRID: AB_2335127); mouse anti-V5, Bio-Rad (MCA1360GA, RRID: AB_567249); mouse anti-CD44, Cell Signaling Technology (3570, RRID: AB_2076465); rabbit anti-EEA1, Thermo Fisher (PA1-063A, RRID: AB_2096819); mouse anti-α tubulin, Sigma-Aldrich (T5168, RRID: AB_477579); rabbit anti-Histone H3, Cell Signaling Technology (4499, RRID: AB_10544537); mouse anti-GFP, Santa Cruz Biotechnology (sc-9996, RRID: AB_627695); mouse anti-mCherry, Novus Biologicals (NBP1-96752, RRID: AB_11034849); rabbit anti-phospho-Threonine/Tyrosine, Cell Signaling Technology (9381, RRID: AB_330301); goat anti-mouse HRP, Jackson ImmunoResearch Labs (115-035-146, RRID: AB_2307392); goat anti-rabbit HRP, Jackson ImmunoResearch Labs (111-035-144, RRID: AB_2307391); donkey anti-mouse Alexa Fluor 488, Invitrogen (A-21202, RRID: AB_141607).

### Other reagents

Lipofectamine 2000, Invitrogen (11668019); Polybrene Infection/Transfection Reagent, Sigma-Aldrich (TR-1003); GFP-Trap magnetic agarose, ChromoTek (gtma-20); Trypsin Gold Mass Spectrometry Grade, Promega (V5280); GFP recombinant protein (Invitrogen, A42613); mCherry recombinant protein (Origene, TP790040); D-biotin, Chem-Impex (00033, Cas #: 58-85-5); (+)-S-Trityl-L-cysteine, Alfa Aesar (L14384, CAS #: 2799-07-7); Puromycin dihydrochloride, Sigma-Aldrich (P8833, CAS # 58-58-2); Doxycycline hyclate, Acros (446061000, CAS #: 24390-14-5); cOmplete Protease Inhibitor Cocktail, Roche (5056489001); Sodium fluoride, Chem-Impex (01523, CAS #: 7681-49-4); Sodium orthovanadate, MP Bio (0215966410, CAS #: 13721-39-6); Oxotremorine M, Santa Cruz Biotechnology (sc-203656, CAS #: 63939-65-1); Phorbol-12-Myristate-13-Acetate, Santa Cruz Biotechnology (sc-3576, CAS #: 16561-29-8); Bryostatin 1, Cayman Chemical (14802, CAS #: 83314-01-6); Gö 6983, Cayman Chemical (13311, CAS #: 133053-19-7); Sotrastaurin, Cayman Chemical (16726, CAS #: 425637-18-9); PF 670462, Fisher Scientific (50-115-0729, CAS #: 950912-80-8); D 4476, Fisher Scientific (2902/10, CAS #: 301836-43-1); Tautomycetin, Tocris (2305, CAS #: 119757-73-2); Fostriecin, VWR (102987-176, CAS #: 87860-39-7); Okadaic acid, Cayman Chemical (10011490, CAS #: 78111-17-); Calyculin A, Santa Cruz Biotechnology (sc-24000, CAS #: 101932-71-2); Phos-tag™ Acrylamide, NARD (AAL-107); 5-gluoro-2-indolyl des-chlorohalopemide (FIPI), Cayman Chemical (13563, CAS #: 939055-18-2) 5-Hexyn-1-ol, Sigma-Aldrich (302015, CAS #: 928-90-5); AZDye 647 Azide, Click Chemistry Tools (1299); 32% Paraformaldehyde, Electron Microscopy Sciences (15714); Streptavidin Alexa Fluor 568, Invitrogen (S11226); Pierce™ High Capacity Streptavidin Agarose, Thermo Fisher (20359); Streptavidin-conjugated HRP, GeneTex (GTX85912); ProLong™ Diamond Antifade Mountant with DAPI, Thermo Fisher (P36971); Clarity™ Western ECL Substrate, Bio-Rad (1705061); BCA Protein Assay Kit, Thermo Fisher (23225); Trizol (Thermo Fisher, 15596026); SMART MMLV reverse transcriptase (Takara, 639524); PerfeCTa SYBR Green SuperMix (Quantabio, 95053). The following compounds were prepared as described previously: 3-azido-1-propanol^66^; BCN-BODIPY^67^.

### siRNA duplexes

| Design ID | Sense |
| --- | --- |
| Negative Control 1 | rCrGrUrUrArArUrCrGrCrGrUrArUrArArUrArCrGrCrGrUAT |
| hs.Ri.CSNK1A1.13.1 | rArCrArUrGrGrArArUrCrArUrUrArGrGrArUrArUrGrUrUTT |
| hs.Ri.CSNK1D.13.1 | rArCrArCrGrCrArCrCrUrUrGrGrArArUrUrGrArArCrArATC |
| hs.Ri.CSNK1E.13.2 | rArUrArUrCrUrArCrCrUrGrGrGrUrGrCrCrArArCrArUrCGC |
| hs.Ri.PRKCA.13.2 | rGrArUrGrArArGrArCrGrArGrCrUrArUrUrUrCrArGrUrCTA |
| hs.Ri.PRKCD.13.1 | rGrGrGrArUrArUrArUrCrArGrGrGrUrUrUrCrGrArGrArAGA |
| hs.Ri.PPP1CA.13.1 | rGrCrArArGrArGrArCrGrCrUrArCrArArCrArUrCrArArACT |
| hs.Ri.PPP1CB.13.1 | rArCrUrArUrUrGrGrArUrGrUrGrArUrUrCrUrArArArArGCT |
| hs.Ri.PPP2CA.13.1 | rArArUrCrArUrGrGrArArCrUrUrGrArCrGrArUrArCrUrCTA |

| Design ID | Antisense |
| --- | --- |
| Negative Control 1 | rArUrArCrGrCrGrUrArUrUrArUrArCrGrCrGrArUrUrArArCrGrArC |
| hs.Ri.CSNK1A1.13.1 | rArArArArCrArUrArUrCrCrUrArArUrGrArUrUrCrCrArUrGrUrCrA |
| hs.Ri.CSNK1D.13.1 | rGrArUrUrGrUrUrCrArArUrUrCrCrArArGrGrUrGrCrGrUrGrUrUrG |
| hs.Ri.CSNK1E.13.2 | rGrCrGrArUrGrUrUrGrGrCrArCrCrCrArGrGrUrArGrArUrArUrCrU |
| hs.Ri.PRKCA.13.2 | rUrArGrArCrUrGrArArArUrArGrCrUrCrGrUrCrUrUrCrArUrCrUrU |
| hs.Ri.PRKCD.13.1 | rUrCrUrUrCrUrCrGrArArArCrCrCrUrGrArUrArUrArUrCrCrCrArA |
| hs.Ri.PPP1CA.13.1 | rArGrUrUrUrGrArUrGrUrUrGrUrArGrCrGrUrCrUrCrUrUrGrCrArC |
| hs.Ri.PPP1CB.13.1 | rArGrCrUrUrUrUrArGrArArUrCrArCrArUrCrCrArArUrArGrUrGrU |
| hs.Ri.PPP2CA.13.1 | rUrArGrArGrUrArUrCrGrUrCrArArGrUrUrCrCrArUrGrArUrUrGrC |

### Quantitative real time PCR primers

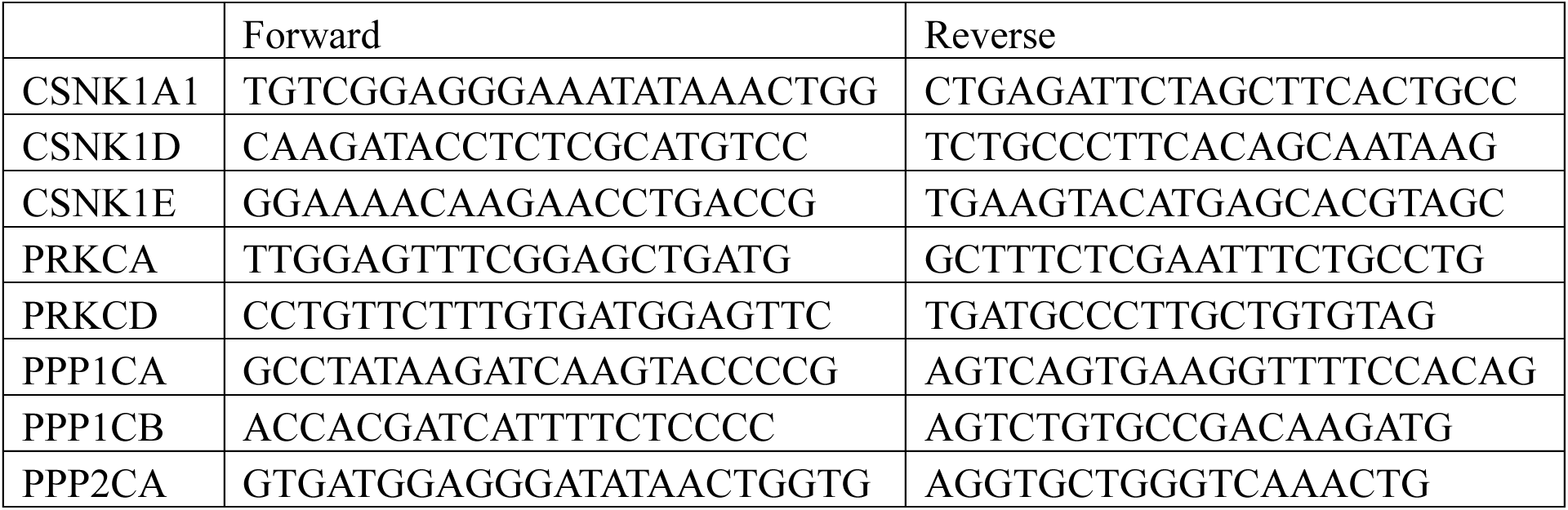

### Plasmids and cloning

EGFP-tagged PLEKHA4 and PLEKHA5 constructs with a CMV promoter were generated in our previous studies^17,20^. mScarlet-i-PLEKHA5^WW^ was made by inserting the cDNA sequence encoding the tandem WW domain (residue 1-105) into the CMV-mScarlet-i-C1 vector (Addgene 85044) using BamHI and SalI. For amino acid mutagenesis, mutations were incorporated into PLEKHA4 and PLEKHA5 plasmids using the QuikChange XL Site-directed mutagenesis kit (Agilent). A nuclear export signal with the amino acid sequence LQLPPLERLTLD was inserted either between PLEKHA5 and GFP or downstream of GFP into the PLEKHA5^H-BP-PH^-EGFP plasmid, and right after mScarlet-i into the CMV-mScarlet-i-N1 vector (A gift from Barbara Baird’s lab, Cornell University) using the QuikChange XL Site-directed mutagenesis kit.

To construct the single MARS plasmid (PLEKHA5^H-BP-PH(AA)^-mSi-NES), the H-BP-PH domain with K163A/R164A mutations was amplified from the GFP fusion construct and inserted into the EcoRI digested mScarlet-i-N1-NES vector. Additionally, PLEKHA5^H-BP-PH(AA)^ -NES-mSi was made by subcloning H-BP-PH(AA) into an EcoRI-digested CMV-mScarlet-i-N1 vector and then inserting the NES between PLEKHA5 and mScarlet-i using the QuikChange XL Site-directed mutagenesis kit. An additional H-BP-PH(AA) motif was added to PLEKHA5^H-BP-PH(AA)^-mSi-NES and PLEKHA5^H-BP-PH(AA)^-NES-mSi at the KpnI site to generate 2xMARS tandem constructs, with the two H-BP-PH(AA) motifs separated by a flexible linker (SGGRGSGSGSGSG). Subsequently, an ORF encoding HA-nGFP (amplified from pcDNA3.1-HA-nGFP-(EAAAK)4-OGT(4), Addgene 160739) was inserted into the PLEKHA5^H-BP-PH(AA)^-mSi-NES plasmid using the BamHI site to yield the MARS-nGFP construct (PLEKHA5^H-BP-PH(AA)^-HA-nGFP-mSi-NES). 2xMARS-nGFP was generated similarly by adding HA-nGFP to the 2xPLEKHA5^H-BP-PH(AA)^-mSi-NES plasmid between the tandem H-BP-PH(AA) motifs and mScarlet-i. MARS (PLEKHA5^H-BP-PH(AA)^-mSi-NES) and 2xMARS-nGFP constructs (2xPLEKHA5^H-BP-PH(AA)^-HA-nGFP-mSi-NES) were then subcloned to a modified pCDH-CMV-MCS-EF1α-Puro vector (with an AscI site inserted upstream of the XbaI site in the original plasmid) by AscI and NotI to yield the lentiviral plasmid.

For nGFP recruitment experiments, PGK-GFP was made by swapping the CMV promoter in CMV-EGFP-C1 (Clontech) vector with the hPGK promoter amplified from pLenti.PGK.blast-Renilla_Luciferase (Addgene 74444) using AseI and NheI. pCDH-CMV-EGFP-FLAG was cloned by inserting the FLAG tag (DYKDDDDK) after EGFP in pCDH-CMV-EGFP^68^ using the QuikChange XL Site-directed mutagenesis kit. pLenti-PGK-EGFP-FLAG was cloned by inserting EGFP-FLAG (amplified from pCDH-CMV-eGFP-FLAG) to SalI and BsrGI-digested pLenti.PGK.blast-Renilla_Luciferase (Addgene 74444).

Single and tandem MARS fusions to WT PLD_PMF_ were made by replacing Lyn_10_ in pCDNA3-Lyn_10_-mCherry-PLD_PMF_ (Julia Li, Baskin Lab) for either H-BP-PH(AA)-NES or 2xH-BP-PH(AA)-NES using HindIII and AgeI. Subsequently, the H170A mutation of PLD was incorporated using the QuikChange XL Site-directed mutagenesis kit into these plasmids. To make MARS-p85^iSH2^ and 2xMARS-p85^iSH2^, the p85^iSH2^ sequence (amplified from pPBpuro-mNG-eDHFR(69K6)-p85iSH2, Addgene 172103) was first inserted after the mCherry in pCDNA3-Lyn_10_-mCherry-PLD_PMF_ by BamHI and ApaI to replace PLD_PMF_, yielding pCDNA3-Lyn_10_-mCherry-p85^iSH2^. Subsequently, the Lyn_10_ tag was again replaced with either H-BP-PH(AA)-NES or 2xH-BP-PH(AA)-NES using HindIII and AgeI. CMV-GFP-PLD_PMF_ and PGK-GFP-p85^iSH2^ were constructed by inserting the coding sequence for each gene into the CMV-EGFP-C1 expression vector (Clontech) and PGK-EGFP-C1 plasmid respectively via HindIII and KpnI.

For TurboID experiments, pCDNA3-Lyn_10_-TurboID-V5 was cloned by replacing the C1(1-29) sequence in C1(1-29)-TurboID-V5-pCDNA3 (Addgene, 107173) with a Lyn_10_ tag (amino acid sequence GCIKSKRKDG). Then the entire Lyn_10_-TurboID-V5 cassette was inserted into the pSBtet-Pur plasmid (Addgene 60507) linearized by NcoI and BspDI to replace the firefly luciferase downstream of the Tet-responsive promoter, generating the pSBtet-Lyn_10_-TurboID-V5-puro plasmid used for making a HeLa cell line stably expressing Lyn_10_-TurboID-V5 under doxycycline induction.

H2B-tagBFP was cloned by inserting the H2B cDNA (amplified from pLenti6-H2B-mCherry, Addgene 89766) into the tagBFP-N1 vector (Evrogen) via SalI and ApaI, and subsequently H2B-tagBFP was cloned into a modified pCDH-CMV-MCS-EF1α-Puro vector (inserted an AscI site upstream of the XbaI site in the original plasmid) by AscI and NotI to yield the lentiviral plasmid. Akt^PH^-miRFP670 was generated by cloning the Akt^PH^ sequence (amplified from GFP-Akt^PH^, a gift from the De Camilli lab at Yale University) into the miRFP670-N1 vector (Addgene 79987) using SalI and ApaI.

### Cell culture

Flp-In T-Rex HeLa cells (Thermo Fisher) were cultured in DMEM (Corning) supplemented with 10% FBS (Corning) and 1% penicillin/streptomycin (P/S, Corning) at 37 °C in a 5% CO_2_ atmosphere. HEK 293TN cells (gift from Anthony Bretscher, Cornell University) were cultured under the same conditions with additional supplement of 1 mM sodium pyruvate (Corning) in the media. HeLa cell line with PLK1 C-terminally tagged by GFP at the endogenous locus via CRISPR/Cas9 was a gift from Prof. Iain Cheeseman (MIT). H2B-mCherry-expressing Flp-In T-Rex HeLa cells were made in our previous study^17^. To generate a stable cell line expressing Lyn_10_-TurboID-V5 under doxycycline induction (2.5 μg/mL), pSBtet-Lyn_10_-TurboID-V5-puro and pCMV(CAT)T7-SB100 (Addgene 34879) were co-transfected into Flp-In T-Rex HeLa cells in a 19:1 ratio, and cells were selected with 2 μg/mL puromycin treatment beginning 24 h after transfection and continuing for 2 days. Stable expression of H2B-tagBFP, MARS, and 2xMARS-nGFP was accomplished by transduction of corresponding lentiviral plasmids into Flp-In T-Rex HeLa cells and a similar puromycin selection protocol. To generate cell lines stably expressing both 2xMARS-nGFP and various levels of GFP for stoichiometry estimation, 2xMARS-nGFP stable cells were transduced with pLenti-PGK-GFP-FLAG or pCDH-CMV-GFP-FLAG, lifted from the 10 cm dish, centrifuged (500 g, 3 min) to remove media, and resuspended in 1 mL FACS buffer (PBS with 0.5% FBS) supplemented with 1% P/S. Cells were then sorted using a Sony MA900 sorter. pLenti-PGK-GFP-FLAG cells were gated on mScarlet-positive cells (90%) then on GFP-positive cells (90%), and 1.5 million cells were collected for grow back. pCDH-CMV-GFP-FLAG cells were gated on mScarlet-positive cells (93%) then split into three fractions based on GFP level. 1.1 million “GFP-low” (bottom 15%), 1.5 million “GFP-medium” (middle 20%), and 1.1 million “GFP-high” (top 15%) cells were collected for grow back. Cells were passaged twice after collection before live cell imaging or Western blot experiments. Cell lines were used without further authentication, and mycoplasma testing (MycoSensor PCR assay, Agilent) was performed yearly.

### Plasmid transfection

Plasmids were transfected into mammalian cells using Lipofectamine 2000 following the manufacturer’s protocol, except that OptiMEM was substituted with Transfectagro. Cells were incubated with the transfection mix in Transfectagro with 10% FBS for 7–8 h and then cultured in regular growth medium until harvest.

### siRNA knockdown

DsiRNA duplexes were obtained from Integrated DNA Technologies. Transfections with siRNA were performed using Lipofectamine RNAiMAX (Thermo Fisher, 13778150) following the manufacturer’s protocol except using Transfectagro in place of OptiMEM. Cells were incubated with transfection mix in Transfectagro supplemented with 10% FBS for 12–16 h, followed by exchange with fresh media. NC1 (negative control 1, IDT) was used as the control siRNA duplex. 24 h post transfection, cells were analyzed via qRT-PCR or Western blot. For qRT-PCR analysis, each siRNA was transfected individually. For Western blot analysis, siRNAs targeting three CK1 isoforms (CSNK1A1, CSNK1D, and CSNK1E), two PKC isoforms (PRKCA AND PRKCD), and three PP1/PP2A catalytic subunits (PPP1CA, PPP1CB, and PPP2CA) were co-transfected together, respectively.

### Lentivirus production

HEK 293TN cells were seeded one day before transfection to achieve approximately 50% confluency on the day of transfection with lentiviral plasmids. Packaging plasmids PAX2 and VSVg (gifts from Lammerding lab, Cornell University), along with the lentiviral plasmid encoding proteins of interest, were co-transfected into HEK 293TN cells in the ratio of 3.1:1:4.2 using Lipofectamine 2000 overnight in Transfectagro. Transfectagro was then replaced with fresh growth medium the next morning (14–16 h post transfection). The virus-containing medium was collected four times every 12 h starting from 24–30 h post transfection, filtered through a 0.45 μm syringe filter, and stored at 4 °C for up to 48 h if not used immediately.

### Lentivirus transduction

Cells were seeded on 6-well plates one day before transduction to achieve around 70% confluency the next day. One “kill control” well was included in the selection process where cells in this well would not be treated with virus medium. During viral transduction, cells in each well were incubated with 1 mL of fresh growth medium, 2 mL of virus-containing medium, and 8 μg/mL of polybrene. The transduction process was repeated three times every 12 h, and after the last transduction, fresh medium was supplemented for 12 h before the selection process. Cells in 6-well plates were trypsinized, seeded into 10-cm dishes, and incubated with puromycin until all the kill control cells died. During the selection, drug-containing media was exchanged every two days. Stable expression of proteins of interest was examined by confocal microscopy or Western blot analysis.

### Cell synchronization

To synchronize cells in prometaphase, 5 μg/mL of S-trityl-L-cysteine (STLC) was added to confluent dishes of cells for 16 h. Mitotic cells were dislodged from dishes by gentle tapping, washed with 1X DPBS, and collected. Cells pellets were stored at −80 °C if not used immediately.

### Confocal microscopy

For live-cell imaging, cells were seeded on 35 mm glass bottom culture dishes (14 mm diameter, #1.5 thickness, Matsunami Glass). For immunofluorescence, cells were seeded on 12 mm cover glass (#1.5 thickness, Fisherbrand) in 12-well plates (Corning). Cells were imaged either live or fixed by immunofluorescence 24–30 h post transfection. Images were acquired via the Zeiss Zen Blue 2.3 software on a Zeiss LSM 800 confocal laser scanning microscope equipped with Plan Apochromat objectives (40x 1.4 NA) and two GaAsP PMT detectors. Solid-state lasers (405, 488, 561, and 640 nm) were used to excite DAPI/tagBFP, EGFP/Alexa Fluor 488, mCherry/ mScarlet-i/Alexa Fluor 568, and miRFP/miRFP670 respectively. Live-cell time-lapse movies were acquired using definite focus. Depending on the experiments, images and movies were acquired either at room temperature or at 37 °C with a 5% CO_2_ atmosphere when indicated. Acquired images were analyzed using FIJI. To quantify PM and cytosolic fluorescence, five line scans at different areas were drawn for each cell to get the average fluorescence. The Coloc 2 command was used to compute the Pearson coefficients with the PLEKHA5 channel set as reference ROI or mask.

### Live-cell imaging

For most live imaging experiments not involving time-lapse movies, cells were transiently transfected with expression plasmids (unless stated otherwise) for 24–30 h, rinsed twice, and then imaged in Tyrode’s-HEPES buffer (T/H buffer: 135 mM NaCl, 5 mM KCl, 1.8 mM CaCl2, 1 mM MgCl2, 5 mg/mL glucose, 5 mg/mL bovine serum albumin, 20 mM HEPES, pH 7.4) at room temperature.

For the PI(4,5)P_2_ depletion assay, three plasmids, PLEKHA5^H-BP-PH^-EGFP, M1R-mCherry, and iRFP-PLCd^PH^, were co-transfected into HeLa cells for 24–30 h. Cells were rinsed with T/H buffer twice and subjected to time-series acquisition with definite focus at room temperature. Images were acquired every 5 s for in 5 min total, and 10 μM oxotremorine-M was added 30 s after starting the acquisition to activate M1R for acute PI(4,5)P_2_ depletion at the plasma membrane. For PKC activation imaging experiments, cells were transfected with GFP-tagged PLEKHA5 constructs for 24–30 h and then treated with PKCi (100 nM Gö 6983 and 100 nM Sotrastaurin/AEB071) or vehicle for 1 h. T/H buffer was used to rinse the cells twice and added to cells with fresh PKCi or vehicle. PKC agonists, PMA (100 nM) or Bryostatin 1 (200 nM), were added to the dishes, and immediately afterward time-lapse acquisition with definite focus was started for 20-25 min at 37 °C with a 5% CO_2_ atmosphere. Images were acquired every 1 min.

For long-term time-lapse imaging, cells were imaged in DMEM lacking phenol red (Gibco) supplemented with 10% FBS and 1% P/S at 37 °C in a 5% CO_2_ environment 24–30 h post transfection. For PLEKHA5^WW-H-BP-PH^-EGFP and H2B-mCherry imaging to track PLEKHA5 PM translocation, images were acquired every 1 min for 12 h. For H2B-tagBFP and MARS or 2xMARS-nGFP imaging to monitor MARS localization, images were acquired every 2 min for 14 h. For H2B-tagBFP, GFP-Akt^PH^, and MARS or MARS-p85^iSH2^ imaging to monitor mitosis progression, images were acquired every 4 min for 14 h.

#### Immunofluorescence

For immunofluorescence analysis of Lyn_10_-TurboID-V5, cells were fixed in 4% paraformaldehyde for 15 min, permeabilized with 0.5% Triton-X 100 in 1X PBS for 5 min, and blocked in 2% BSA and 0.1% Tween-20 in 1X PBS (blocking buffer) for 30 min. Cells were treated with primary antibody (anti-V5) in blocking buffer for 1 h, rinsed with wash buffer three times and incubated with secondary antibodies in blocking buffer for 1 h. Cells were then rinsed with wash buffer three times and with 1X PBS twice, mounted on slides in ProLong Diamond Antifade with DAPI, and incubated overnight in dark before imaging. All steps were performed at room temperature, and slides were stored at 4 °C.

### Proximity biotinylation with Lyn_10_-TurboID

pSBtet-Lyn_10_-TurboID-V5 expression were induced in HeLa cells with 2.5 μg/mL doxycycline for 48 h and then incubated with 500 μM biotin for 10 min at 37 °C under 5% CO_2_. For localization analysis by immunofluorescence, cells were rinsed five times with 1X PBS and then prepared as described earlier (see Confocal microscopy section). For streptavidin-agarose pulldown and Western blot, cells were rinsed five times with 1X PBS and lysed with RIPA buffer (150 mM NaCl, 25 mM Tris pH 8.0, 1 mM EDTA, 0.5% sodium deoxycholate, 0.1% SDS, and 1% Triton X-100) supplemented with cOmplete protease inhibitor cocktail. Cell lysates were sonicated with four pulses at 20% intensity using a tip sonicator and centrifuged at 13,000 *g* for 5 min at 4 °C to clarify lysates. Protein concentration was quantified using the BCA assay (Thermo Fisher), and a small fraction of the clarified cell lysates was saved and normalized as input. The remaining cell lysate was subjected to pulldown using streptavidin-agarose with rotation at 4 °C overnight (12-16 h). The resin was then centrifuged for 5 min at 1000 *g*, washed two times with RIPA buffer, one time with 1 M KCl, one time with 0.1 M Na_2_CO_3_, one time with 2 M urea in 10 mM Tris pH 8.0, and two times with RIPA buffer to reduce non-specific binding. Samples were then denatured and analyzed by SDS-PAGE and Western blot.

### PI(4,5)P_2_ depletion and analysis

A 60 mm dish was seeded with 400,000 HeLa cells and incubated overnight. The dish was then transfected with M1R-mCherry with 2000 ng plasmid and 4 µL Lipofectamine 2000. 6 h after the transfection, the DNA-containing medium was replaced with fresh growth medium, and the dish was incubated for another 24 h. Subsequently, the dish was rinsed with PBS twice and treated with 10 µM oxotremorine-M in Tyrode’s HEPES buffer for 30 s. After the treatment, the dish was immediately rinsed with ice-cold PBS twice and then added 400 µL of ice-cold 2 mM AlCl_3_ in methanol:12N HCl (96:4). The cell monolayer was scraped off, and the resulting suspension was transferred to a 1.5 mL Eppendorf tube, to which was subsequently added 300 µL of chloroform followed by a 1 min vortex. The mixture was added 150 µL ice-cold water and centrifuged at 9600 x g, 4 °C for 2 min. The lower layer (organic layer) was transferred to a new Eppendorf tube containing 1 mL of ice-cold in methanol:2 M oxalic acid (1:0.9). The resulting mixture was vortexed for 1 min and centrifuged at 9600 x g, 4 °C for 2 min. The lower layer (organic layer) was transferred to a new Eppendorf tube and dried under a stream of nitrogen to obtain a lipid thin film. The extracted lipids were deacylated by incubation with 500 µL of 40% methylamine:water:n-butanol:methanol (36:8:9:47) at 50 °C for 45 min. The mixture was dried by speed-vac, resuspended in 500 µL of n-butanol:petroleum ether:ethyl formate (20:40:1) and 500 µL water, and vortexed for 1 min. After centrifugation at 9600 x g, 4 °C for 2 min, the lower layer (aqueous layer that contained the deacylated products) was transferred to a new Eppendorf tube. The organic phase was extracted with additional 500 µL water, and the aqueous layer was combined and dried by speed-vac overnight with no heat. Samples were resuspended in water for detection on the Integron High Performance Ion Chromatography (HPIC) with suppressed conductivity detection (Thermo Fisher) as described previously^69,70^.

### Western blot

Except for the Lyn_10_-TurboID experiments and the 2xMARS-nGFP and GFP stoichiometry experiments, HeLa cells were transfected with expression plasmids for Western blot analysis. 48 h post transfection, cells were lysed with RIPA buffer (150 mM NaCl, 25 mM Tris pH 8.0, 1 mM EDTA, 1% Triton X-100, 0.5% sodium deoxycholate, and 0.1% SDS) supplemented with cOmplete protease inhibitor cocktail and a phosphatase inhibitors mixture (20 mM sodium fluoride, and 1 mM sodium orthovanadate) on ice. Cell lysates were sonicated for 2 pulses at 20% intensity and centrifuged at 13,000 *g* for 5 min at 4 °C. Protein concentrations of the clarified cell lysates were quantified using the BCA assay. For Phos-tag SDS-PAGE, ZnCl_2_ solution was added to cell lysates to reach a final Zn^2+^ concentration of 1 mM to counteract the EDTA in RIPA buffer, which would interfere with electrophoresis of Phos-tag gels. Cell lysates were normalized to 2–3 mg/mL, and denatured with 6X Laemmli Buffer (13.3% SDS, 0.067% bromophenol blue, 52.2% glycerol, 67 mM Tris pH 6.8 and 11.1% β-mercaptoethanol) at 95 °C for 5 min. Samples were analyzed by Phos-tag or regular SDS-PAGE and Western blot with detection by chemiluminescence using Clarity Western ECL substrate (Bio-Rad). Western blots were imaged using a ChemiDoc MP Imaging System (Bio-Rad) and quantified with Image Lab software (Bio-Rad).

### 2xMARS-nGFP and GFP stoichiometry estimation

Whole cell lysates harvested from HeLa cells stably expressing 2xMARS-nGFP-mSi and PGK-GFP, CMV-GFP low, medium, and high were normalized, and 20 µg of total protein was analyzed by Western blot. Using the 4PL model in GraphPad Prism, standard curves of purified recombinant GFP and mCherry proteins were generated and the mass of GFP and 2xMARS-nGFP-mSi in cell lysates were interpolated. The quantity of these proteins in moles was calculated by dividing the mass amount by the molecular weight, and the molar ratios of the GFP to 2xMARS-nGFP were plotted as an estimation of stoichiometry. For live-cell imaging, these stable cells were seeded on 35 mm glass bottom culture dishes 24 h before image acquisition.

### Cell proliferation assay

2,000 HeLa cells, either untransduced or transduced with lentivirus encoding MARS or 2xMARS-nGFP, were seeded in each well of a low-evaporation 96-well plate (Corning), two wells per cell line. Three images per well were acquired every 15 min for 96-120 h in an IncuCyte incubator with a 20× objective. Cell confluency at each time point was analyzed using the IncuCyte ZOOM system. Confluency data within the exponential growth phase were plotted and fitted for doubling time calculation using the Exponential growth curve model in GraphPad Prism.

### Anti-GFP immunoprecipitation

Cells transfected with GFP fusion proteins were harvested and lysed in IP lysis buffer (150 mM NaCl, 1% NP-40, 0.25% sodium deoxycholate, 5 mM EDTA, 50 mM Tris pH 7.5) supplemented with cOmplete protease inhibitor cocktail on ice. Cell lysates were sonicated for 4 pulses at 20% intensity and centrifuged at 13,000 *g* for 5 min at 4 °C. Protein concentration of the clarified cell lysates was quantified using the BCA assay, and a small fraction of the clarified cell lysates was separated and normalized as input. The remaining cell lysate was subjected to immunoprecipitation using GFP-Trap Magnetic Agarose with rotation at 4 °C overnight (12–16 h). The magnetic resin was then separated by a magnetic tube rack, washed three times with lysis buffer (at least 10 times the bead volume), denatured and analyzed by SDS-PAGE and Western blot with detection by chemiluminescence using Clarity Western ECL substrate, or by mass spectrometry-based phosphoproteomics analysis, as described in detail in the next section. Cells harvested for phosphoproteomics analysis were treated with the pan PPP inhibitor Calyculin A (50 nM for 15 min) to enrich phosphorylated proteins.

### Mass spectrometry analysis of phosphopeptides

The GFP-Trap magnetic resin was washed four times with IP lysis buffer and incubated with elution buffer (100 mM Tris pH 8.0, 1% SDS) at 65 °C for 15 min. The collected supernatant was reduced with 10 mM DTT at 65 °C for 10 min and then alkylated with 25 mM iodoacetamide (in 50 mM Tris pH 8.0) at room temperature for 15 min in dark. Proteins from the mixed sample were precipitated in a solution containing 50% acetone, 49.9% ethanol and 0.1% acetic acid (PPT solution) in the −20 °C freezer for 1 h and pelleted at 17,000 *g* for 10 min. The protein pellet was washed with PPT solution once, pelleted again, air-dried, and resuspended in urea/Tris solution (8 M urea, 50 mM Tris pH 8.0) and NaCl/Tris solution (150 mM NaCl, 50 mM Tris pH 8.0) in a ratio of 1:3 respectively, for a final concentration of 2 M urea. One μg of mass spectrometry grade trypsin (Trypsin Gold, Promega; 1 mg/mL in 0.03% acetic acid) was added, and proteins were trypsinized at 37 °C overnight on a nutator.

Protein digests were acidified with 0.25% formic acid (FA) and 0.25% trifluoroacetic acid (TFA), desalted using the Sep-Pak C18 resin (Waters; WAT054945), and dried. The sample was then suspended in 70% acetonitrile with 1% formic acid and subjected to hydrophilic interaction chromatography (HILIC) prefractionation using a high-performance liquid chromatography (HPLC) UltiMate3000 system (Thermo Fisher Scientific). Fractions were then analyzed via nanoLC-MS/MS using an UltiMate 3000 system coupled to a Q Exactive HF mass spectrometer (Thermo Fisher Scientific). Reverse phase chromatography was conducted using a C18 resin (Reprosil Pur C18AQ 3 μm) packed in-house into a 35 cm long column (100 μm inner diameter) using a 70-minute linear gradient from 8% to 30% acetonitrile with 0.1% FA at a flow rate of 300 nl/min. The Xcalibur 4.2.47 software (Thermo Fisher Scientific) was used for data-dependent acquisition, with a scan range of 380 to 1800 m/z and a mass resolution of 60,000. The automatic gain control target was set to 3e6 and the maximum injection time to 30 ms. MS^2^ was performed by the acquisition of the 20 most abundant ions per cycle fragmented with a normalized collision energy of 28. The isolation window for the fragmented ions was set to 2 m/z with a mass resolution of 15,000, a maximum injection time of 100 ms, and an automatic gain control target of 1e5.

For data analysis and identification of phosphopeptides, raw data was processed through the Trans Proteomic Pipeline^71^ package (ver. 6.0.0) with spectral data files searched through the Comet search engine^72^ over the human UniProt proteome database downloaded on 2019-11-06. Searching parameters were set to a mass tolerance of 10 ppm, a semi-tryptic digest requirement with 2 allowed miscleavages, dynamic mass modification of 79.966331 Da for phosphorylated serines, threonines, and tyrosines, and a static modification of 57.021464 Da for alkylated cysteines. Phosphorylation site localization probabilities were determined using PTMProphet. The mass spectrometry proteomics data have been deposited to the ProteomeXchange Consortium via the PRIDE partner repository with the dataset identifier PXD052237^73^.

### Phos-tag SDS-PAGE

#### Phos-tag gel preparation

Phos-tag gels were prepared as described^74^ with slight modification. Briefly, to make 8 mL of 9% resolving Phos-tag gel solution (enough for one 1.5 mm Bio-Rad mini gel cast), the following reagents were mixed together evenly: 3.28 mL of deionized water, 2 mL of 1.4 M Bis-Tris·HCl (pH 6.8), 40 μL of 5 mM Phos-tag solution, 40 μL of 10 mM ZnCl_2_ solution, 2.4 mL of 30% acrylamide/0.8% bis-acrylamide, 240 uL 10% APS (w/v), and 8 uL of TEMED. 3 mL of stacking gel solution (enough for one 1.5 mm Bio-Rad mini gel cast) was prepared by mixing together: 1.76 mL of deionized water, 0.75 mL of 1.4 M Bis-Tris·HCl (pH 6.8), 400 uL of 30% acrylamide/0.8% bis-acrylamide, 90 uL of 10% APS (w/v), and 3 uL of TEMED.

#### Phos-tag gel electrophoresis

Running buffer (0.1 M Tris, 0.1 M MOPS, 5 mM sodium bisulfite, and 0.1% SDS, pH 7.8) for Phos-tag gel electrophoresis was prepared fresh every time. Gels were run under constant current (30 mA per gel) for 3 h and then rocked in methanol-free transfer buffer (25 mM Tris·HCl, 96 mM glycine, pH 8.2; 50 mL per gel) plus 1 mM EDTA (pH 8.0) twice for 10 min each to remove Zn^2+^ ions, and once in methanol-free transfer buffer for 10 min before proteins were transferred to nitrocellulose membranes.

### Kinase and phosphatase agonists and inhibitors treatment summary

The following compounds were added to cells in conditions as summarized:

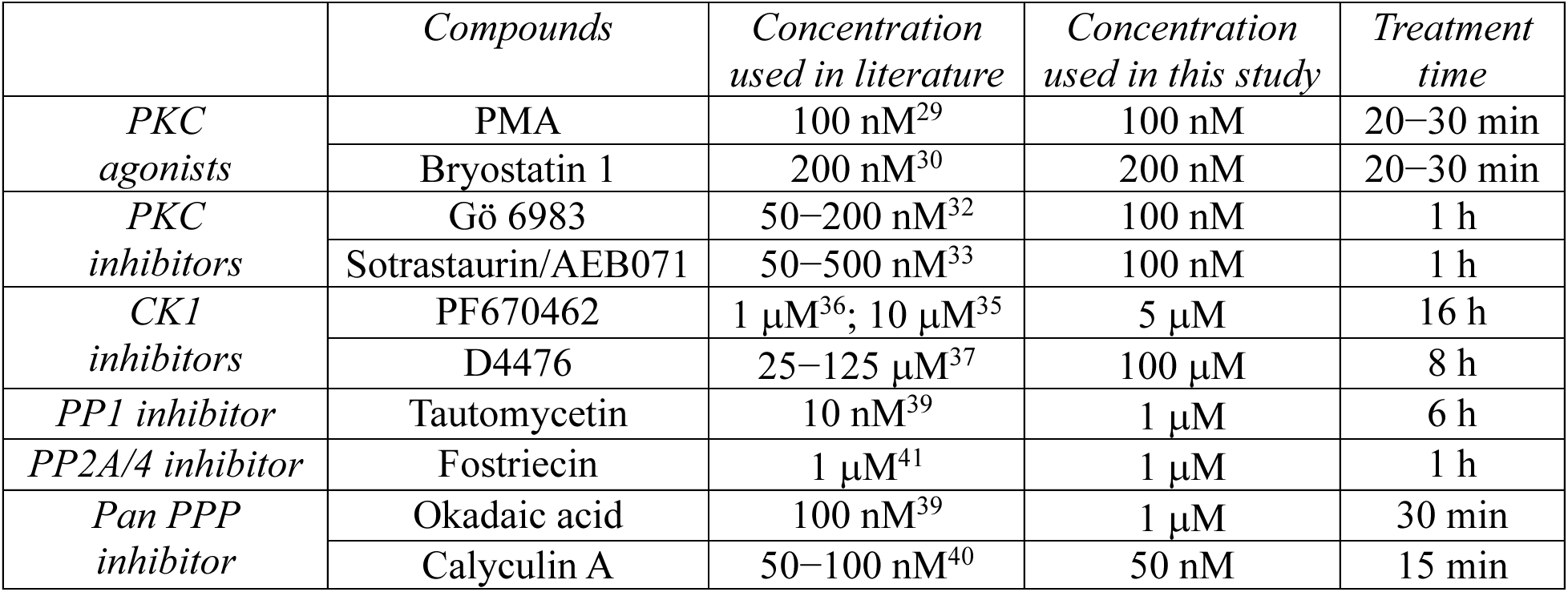

Gö 6983 and Sotrastaurin/AEB071 combined were used as pan-PKC inhibitor (PKCi).

### Quantitative real-time PCR (qRT-PCR)

Cells were seeded on 6-well plates to achieve ∼50% confluency the next day. Cells in each well were then transfected with siRNA duplexes. The following 24 h post transfection, cells were washed with ice-cold DPBS twice and homogenized in 1 mL Trizol (Thermo Fisher). Total RNA was then extracted following the manufacturer’s protocol. 1 µg of total RNA was reverse transcribed to cDNA using SMART MMLV reverse transcriptase (Takara) and Oligo(dT)_20_ as a primer. qRT-PCR was performed on a Light Cycler 480 instrument (Roche). Each PCR reaction contained 0.6 µM of the primers specific to the genes of interest, 15–20 ng of template cDNA and 1X PerfeCTa SYBR Green SuperMix (Quantabio). The program was run for 10 min at 95 °C followed by 45 cycles of heating at 95 °C for 15 s and at 55 °C for 60 s. The data was analyzed using ΔΔC_t_ method.

### IMPACT labeling to measure PLD_PMF_ activity

#### Azidopropanol with BCN-BODIPY labeling for 2xMARS-PLD_PMF_

Cells were transfected with 2xMARS-PLD_PMF_ constructs for 32 h and then treated with 5 µg/mL STLC for 16 h to synchronize cells to prometaphase. The same volume of DMSO was added to asynchronous populations. Cells were either incubated with the PLD inhibitor FIPI (750 nM) or vehicle for 30 min and then treated with 1 mM azidopropanol for 30 min to generate azido phosphatidyl alcohol lipid reporters of PLD activity. Cells were rinsed with 1x PBS three times, followed by incubation with 1 µM BCN-BODIPY for 30 min to fluorescently tag the reporter lipids via strain-promoted azide-alkyne cycloaddition. Cells were then rinsed three times with PBS and incubated in T/H buffer (135 mM NaCl, 5 mM KCl, 1.8 mM CaCl_2_, 1 mM MgCl_2_, 5 mg/mL glucose, 5 mg/mL bovine serum albumin, 20 mM HEPES, pH 7.4) for 15 min to rinse out unreacted, excess BCN-BODIPY. All the incubation steps were performed at 37 °C with a 5% CO_2_ atmosphere. Cells were then washed with PBS twice. Asynchronous cells were lifted following treatment with trypsin-EDTA (Corning; 0.05% trypsin and 0.1% EDTA), and prometaphase cells were dislodged by tapping plates and collected by vigorous pipetting in T/H buffer. Cells were washed with T/H buffer twice, fixed with 4% paraformaldehyde in PBS for 15 min in the dark at room temperature, rinsed twice with FACS buffer (0.1% FBS in PBS), and subjected to two-color flow cytometry analysis. A 561 nm laser was used to excite the mCherry within 2xMARS-PLD_PMF_ constructs, and a 488 nm laser was used to excite BODIPY-derived IMPACT fluorescence as a measure of PLD activity. IMPACT fluorescence from cells with similar expression levels of 2xMARS-PLD_PMF_ constructs were extracted for data analysis. For each experimental condition, IMPACT fluorescence from FIPI-treated samples was averaged and subtracted from each replicate, and the differences of mean fluorescence were plotted.

#### Hexynol with AZDye 647 labeling for 2xMARS-nGFP and GFP-PLD_PMF_

Cells were co-transfected with 2xMARS-nGFP and GFP-PLD_PMF_ constructs for 32 h and then treated with 5 µg/mL STLC for 16 h to synchronize cells to prometaphase. The same volume of DMSO was added to asynchronous populations. Cells were either incubated with the PLD inhibitor FIPI (750 nM) or vehicle for 30 min and then treated with 1 mM hexynol for 30 min to generate alkyne-containing phospholipid reporters of PLD activity. Cells were rinsed with PBS three times. Incubations were performed at 37 °C with a 5% CO_2_ atmosphere. Asynchronous cells were lifted following treatment with 0.05% trypsin-EDTA, and prometaphase cells were dislodged by tapping plates and collected by vigorous pipetting in T/H buffer. Cells were rinsed twice with T/H buffer, fixed with 4% paraformaldehyde in PBS for 15 min in the dark, and rinsed another two times with T/H buffer. Cells were then incubated with the AZDye 647-azide solution (100 mM Tris pH 8, 1 mM CuSO_4_, 50 mM freshly made sodium ascorbate, and 5 μM AZDye 647-azide) for 1 h at room temperature to fluorescently tag the alkyne reporter lipids via Cu-catalyzed azide-alkyne cycloaddition. After being washed with FACS buffer twice, cells were subjected to three-color flow cytometry analysis. A 561 nm laser was used to excite the mScarlet-i within 2xMARS-nGFP, a 488 nM laser was used to excite the GFP-PLD_PMF_ constructs, and a 640 nm laser was used to excite the AZdye 647-derived IMPACT fluorescence as a measure of PLD activity. IMPACT fluorescence from cells with similar expression levels of 2xMARS-nGFP and GFP-PLD_PMF_ were extracted for data analysis. For each experimental condition, IMPACT fluorescence from FIPI-treated samples was averaged and subtracted from each replicate, and the differences of mean fluorescence were plotted.

### Statistical analysis

For all experiments involving quantification, statistical significance was calculated in GraphPad Prism using tests as indicated in the figure legend, including an unpaired two-tailed t-test with unequal variance, a Mann-Whitney U test, or a one-way ANOVA with Tukey or Sidak post hoc multiple comparisons test. The exact *P* values for each comparison are reported, and the number of biological replicates or cells analyzed is stated in the legend. All the raw data were plotted into graphs using GraphPad Prism. For all scatter plots, the black line indicates the mean, and each dot represents an individual biological replicate.

## DATA AVAILABILITY

MARS related plasmids generated in this study have been deposited in Addgene (Deposit 82984). Other data supporting the findings of this study are available within the Article and the Supplementary Information. Source data are provided with this paper.

## ACKNOWLEDGMENTS

We acknowledge support from the NIH (J.M.B.: R01GM143367; S.H.: T32GM138826; M.B.S.: R35GM141159), the Sloan Foundation (J.M.B.: Sloan Research Fellowship), and the DoD (L.B.M.: DURIP GRANT13369767/GRANT13710486). We thank Iain Cheeseman for valuable discussions and for providing the PLK1-GFP cell line, and we thank Lewis Cantley for sharing the kinase predictions of the PLEKHA5 S161 site. We thank Reika Tei, Julia Li, and Neal Vasireddi for technical assistance, the Bretscher lab for the HEK 293TN cell line, the Lammerding lab for the pCDH-CMV-MCS-EF1α-Puro, PAX2 and VSVg plasmids, and access to the IncuCyte system, the Fromme and Emr labs for equipment, and members of the Baskin lab for helpful discussions.

## AUTHOR CONTRIBUTIONS

Conceptualization: X.C., J.M.B.; Funding Acquisition: L.B.M., M.B.S., J.M.B.; Investigation: X.C., S.H., M.M.W., Y.C., D.C., K.M.W., A.L.; Project Administration: L.B.M., M.B.S., J.M.B.; Supervision: L.B.M., M.B.S., J.M.B.; Writing – original draft: X.C., J.M.B.; Writing – review & editing: X.C., S.H., M.M.W., Y.C., D.C., L.B.M., M.B.S., J.M.B.

---

**Extended Data Figure 1. (related to Figure 1).**
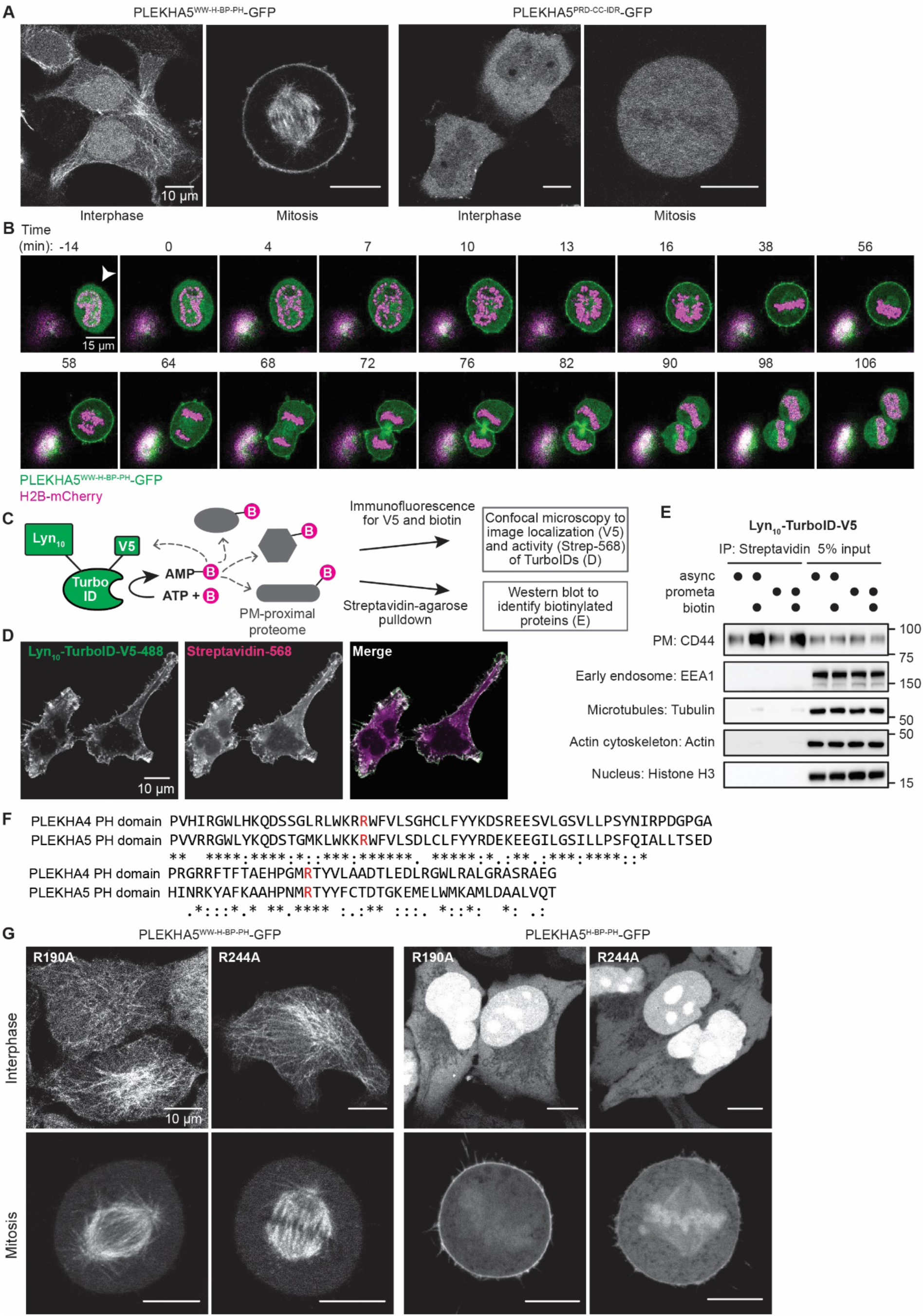
A pool of PLEKHA5 translocates to the PM during mitosis by binding to PI(4,5)P_2_. (A) Live-cell confocal microscopy of HeLa cells transiently expressing GFP-tagged PLEKHA5 constructs in interphase and mitosis. (B) Representative series of still images from a time-lapse movie in asynchronous, stable H2B-mCherry-expressing (magenta) HeLa cells transfected with PLEKHA5^WW-H-BP-PH^-GFP (green). Numbers indicate time before/after nuclear envelope breakdown (defined as t=0 min). (C) Schematic representation of proximity biotinylation strategy using Lyn_10_-TurboID-V5 to biotinylate PM-proximal proteins. (D) Lyn_10_-TurboID showed expected PM-specific localization and biotinylation activity as demonstrated by staining for V5 (green) and biotin (streptavidin-Alexa Fluor 568, magenta). (E) Western blot analysis confirmed that Lyn_10_-TurboID biotinylates a protein marker of the PM but not markers of other subcellular localizations. (F) Clustal Omega alignment of the primary amino acid sequences of the PH domain of PLEKHA4 and PLEKHA5. Highlighted in red are two arginine residues previously established to be required for binding PI(4,5)P_2_ in PLEKHA4. (G) Confocal microscopy of HeLa cells transfected with the R190A and R244A mutants of PLEKHA5^WW-H-BP-PH^-GFP and PLEKHA5^H-BP-PH^-GFP.

**Extended Data Figure 2. (related to Figure 2).**
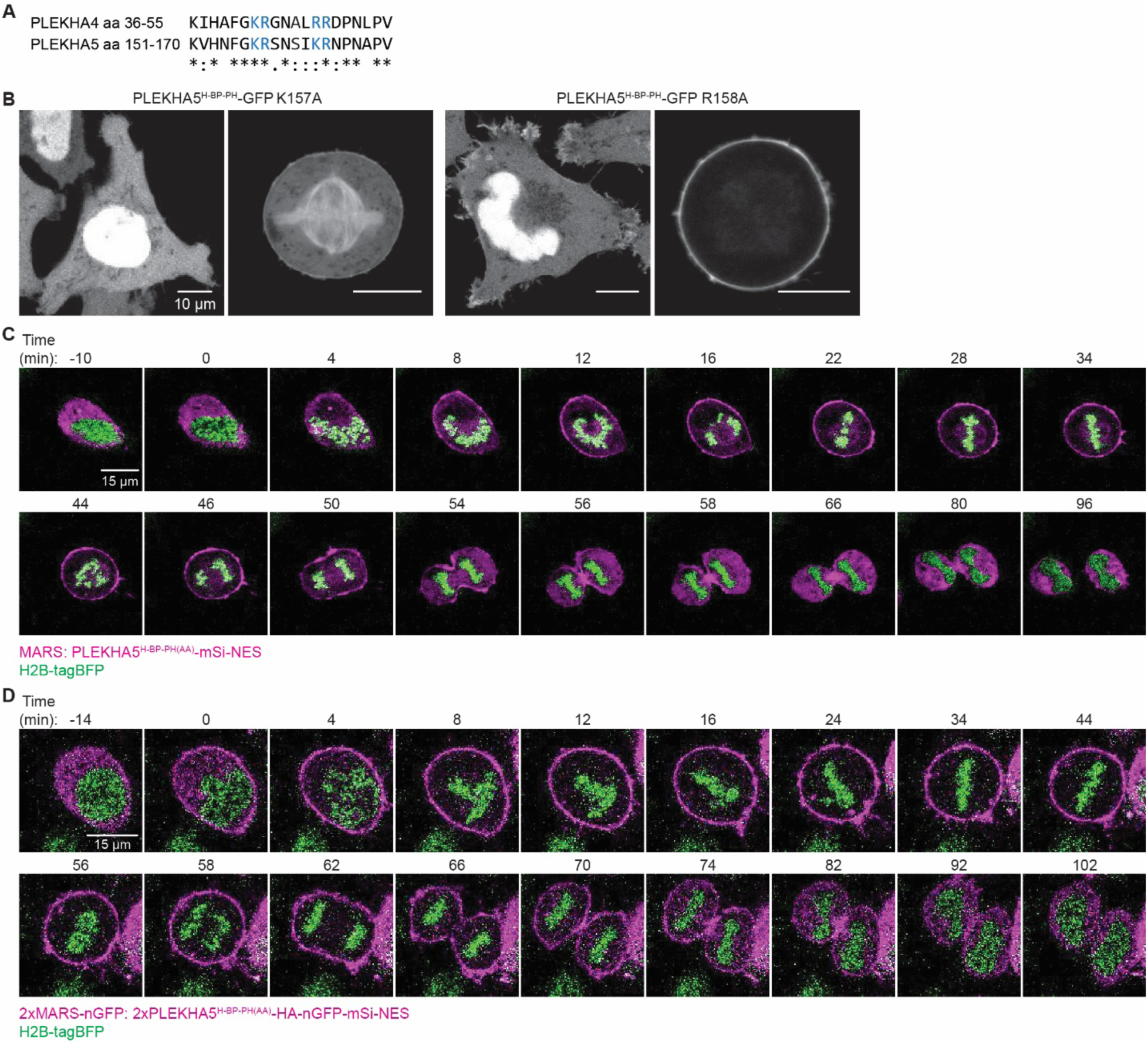
Design and engineering of MARS and MARS-nGFP. (A) Clustal Omega alignment of primary amino acid sequences around the “4A” region of PLEKHA4 and PLEKHA5. (B) Confocal microscopy of the K157A and R158A mutants of PLEKHA5^H-BP-PH^-GFP. (C-D) Representative series of still images from time-lapse movies in asynchronous HeLa cells stably expressing H2B-tagBFP and transfected with either MARS (C) or 2xMARS-nGFP (D). MARS constructs are colored magenta and H2B-tagBFP is colored green. Numbers indicate time before/after nuclear envelope breakdown (defined as t=0 min).

**Extended Data Figure 3. (related to Figure 2).**
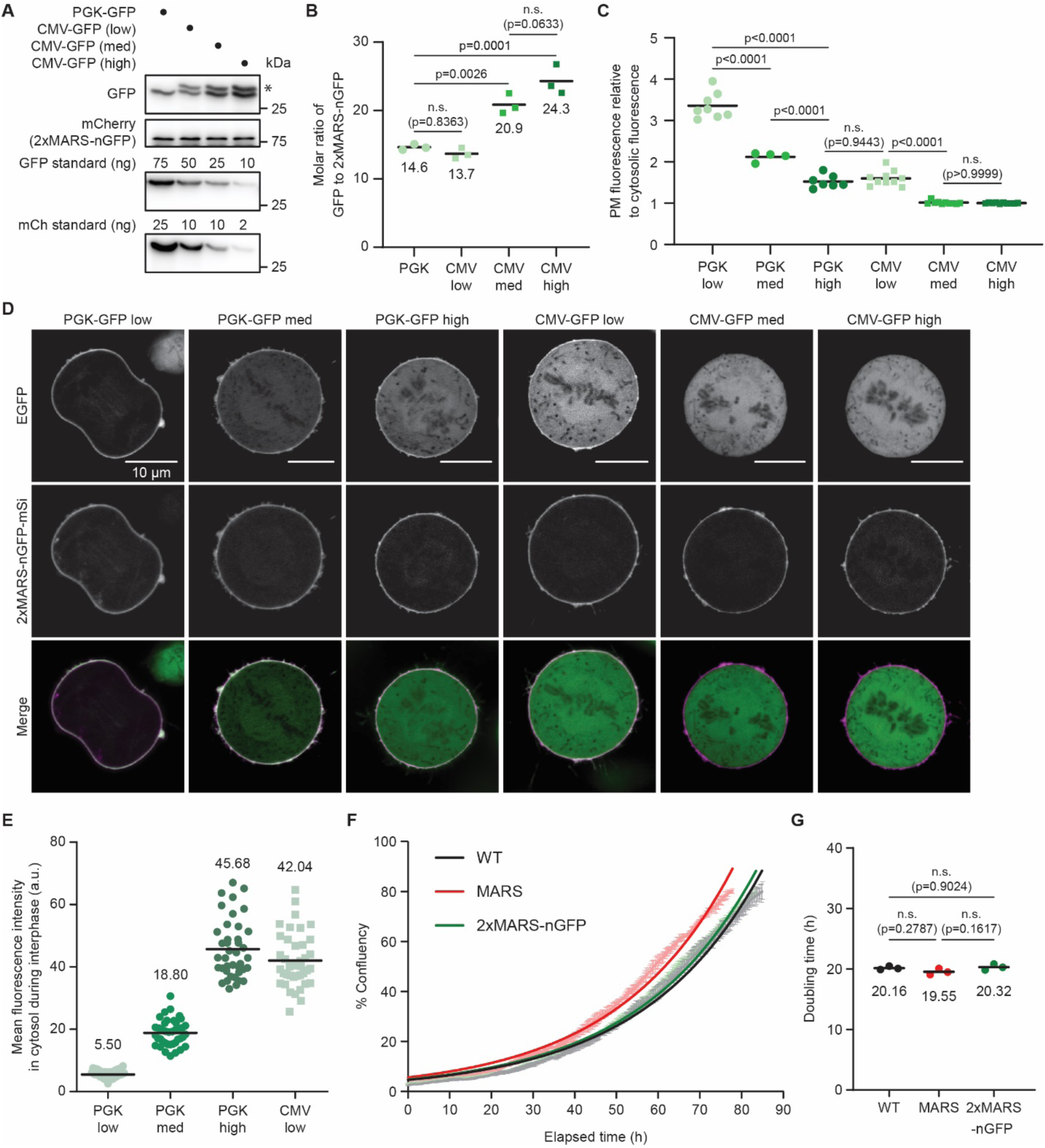
Characterization of stoichiometry of 2xMARS-nGFP and effects of MARS systems on cell proliferation rates. (A) Four stable cell lines were generated via lentiviral transduction that express similar levels of 2xMARS-nGFP and varying levels of GFP, which was under control of either a PGK or CMV promoter (the three CMV-GFP-expressing lines were isolated by FACS, sorting for different levels of GFP expression). Shown are representative Western blots showing GFP (asterisk denotes a longer CMV-GFP product potentially arising from an upstream alternative start codon) and 2xMARS-nGFP (mCherry blot) expression levels in these cell lines alongside blots of purified GFP and mCherry protein standards. (B) Quantification of GFP to 2xMARS-nGFP molar ratios in HeLa cells stably expressing 2xMARS-nGFP and various levels of GFP from (A). (C-D) Imaging analysis of GFP recruitment to the PM during mitosis at different stoichiometries of GFP to 2xMARS-nGFP determined by quantitative image analysis. (C) Quantification of the PM to cytosolic fluorescence ratios of GFP as a readout for measuring GFP recruitment levels by 2xMARS-nGFP. For PGK-GFP: n=8 cells for low GFP expression, n=4 cells for medium expression, and n=7 cells for high expression. For CMV-GFP cell lines, n=10 cells each for low, medium, and high GFP expression level. Cells analyzed came from two independent experiments (one-way ANOVA with Tukey post hoc multiple comparisons test). Note: CMV-GFP cells were sorted by FACS to generate three separate cell lines with low, medium, and high GFP expression, and images were acquired for each cell line. For PGK-GFP cells, the low, medium, and high cells were all from a single cell line and are categorized based on GFP fluorescence intensity during imaging analysis by ImageJ post-acquisition. (D) Representative micrographs for HeLa cells stably expressing 2xMARS-nGFP and varying levels of GFP analyzed in (C). Brightness of the GFP channel was adjusted post-acquisition for better visualization of the weaker fluorescence signals for PGK-GFP and CMV-GFP low cells. (E) Quantification of mean fluorescence intensity in the cytosol during interphase as a means to determine GFP expression levels in PGK-GFP low, medium, high cells and CMV-GFP low cells (n = 40 cells from two independent experiments). Compared to CMV-GFP low cells, PGK-GFP low cells exhibited an average of 7.6x lower expression, PGK-GFP medium cells exhibited ∼2.2x lower expression, and PGK-GFP-high cells exhibited ∼1.1x higher expression. Converting to GFP : 2xMARS-nGFP molar ratio in these cells, the estimated ratios are 1.8 (low), 6.2 (medium), and 15.1 (high). (F-G) Ectopic expression of MARS and 2xMARS-nGFP did not negatively impact rates of cell division and proliferation. (F) Representative growth curves for wild-type (WT) HeLa cells and those stably expressing MARS and 2xMARS-nGFP (n=2 technical replicates for each curve). Raw data (faded-color curves with error bar) acquired by IncuCyte Live-Cell Analysis System were fitted using GraphPad Prism (Exponential growth curve model). Fitted curves are shown as solid lines. (G) Doubling time of the three cell lines calculated from the fitted growth curves (n = 3 biological replicates, one-way ANOVA with Tukey post hoc multiple comparisons test).

**Extended Data Figure 4. (related to Figure 3).**
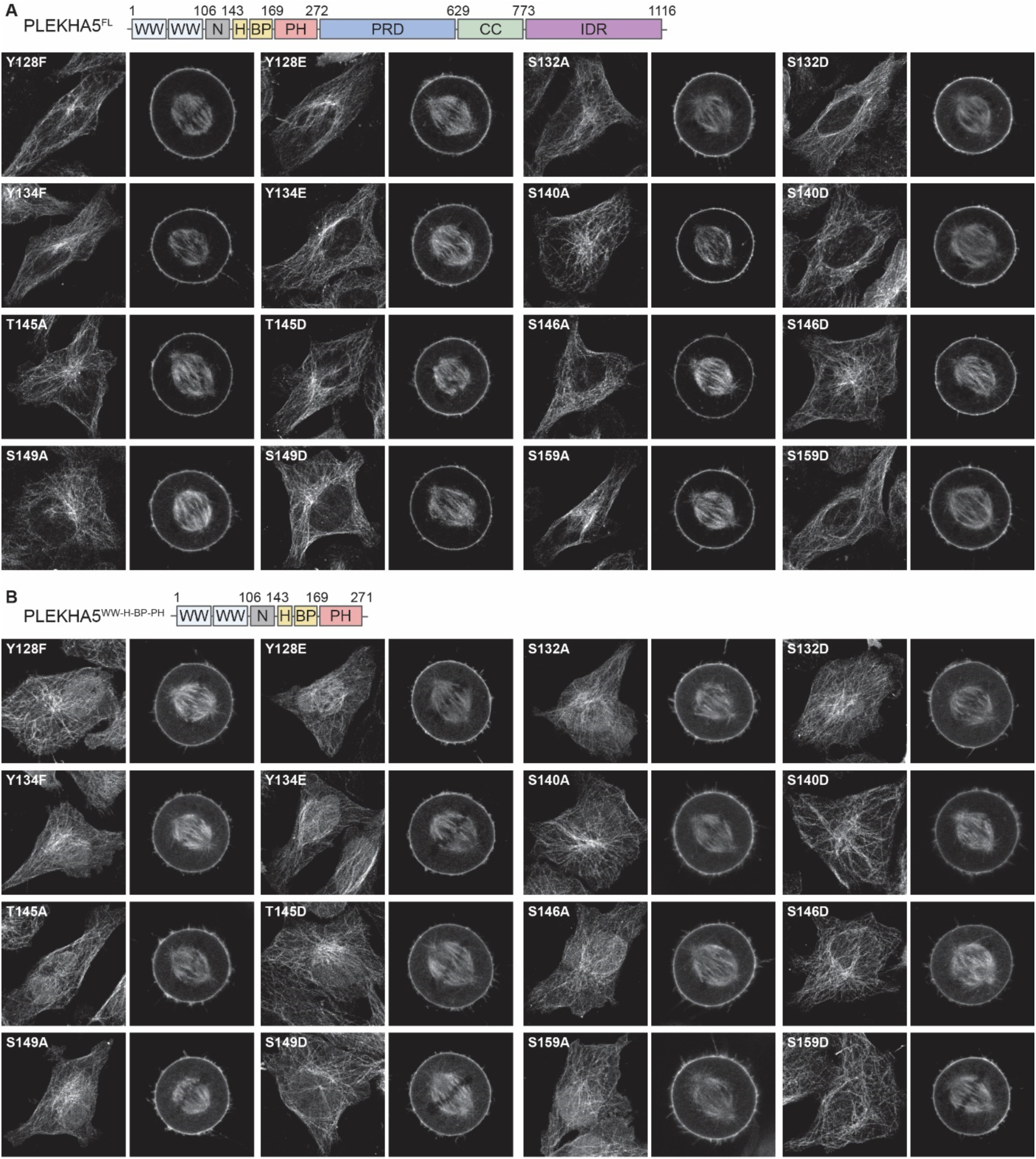
Mutational analysis reveals no contributions of phosphorylation of residues other than S161 to the cell cycle-dependent localization of PLEKHA5. Live-cell confocal microscopy of phosphodeficient (Ser/Thr to Ala and Tyr to Phe) and phosphomimetic (Ser/Thr to Asp and Tyr to Glu) mutants of GFP-tagged PLEKHA5 full-length protein (A) or WW-H-BP-PH fragment (B).

**Extended Data Figure 5. (related to Figure 3).**
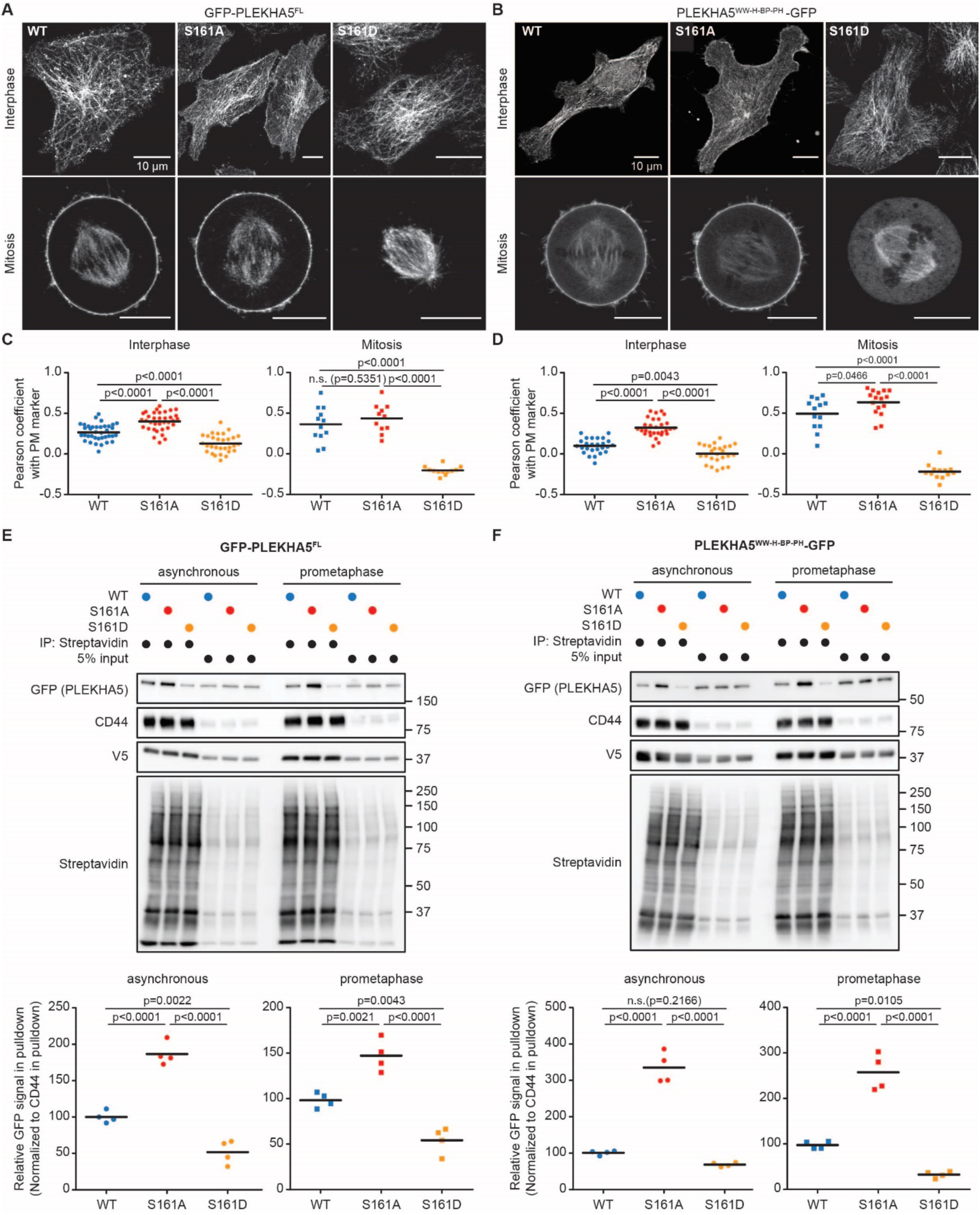
S161 phosphorylation status influences the PM localization of EGFP-PLEKHA5^FL^ and PLEKHA5^WW-H-BP-PH^-EGFP. (A-B) Live-cell confocal microscopy of the subcellular localizations of the WT, S161A, and S161D variants of full-length PLEKHA5^FL^ (A) and the WW-H-BP-PH motifs (B). (C-D) Quantification of PM localization for GFP-PLEKHA5^FL^ (C) and PLEKHA5^WW-H-BP-PH^-GFP (D) using colocalization analysis. Pearson coefficients were computed from images acquired in HeLa cells transiently expressing either of the PLEKHA5 constructs with a transfectable PM marker, Lyn_10_-LOV-mCherry. For GFP-PLEKHA5^FL^ in interphase: n=37 cells for WT, n=35 cells for S161A, and n=30 cells for S161D; in mitosis: n=12 cells for WT and S161D and n=11 cells for S161A. For PLEKHA5^WW-H-BP-PH^-GFP in interphase, n=26 cells for WT, n=29 cells for S161A, and n=25 cells for S161D; in mitosis: n=13 cells for WT and S161D and n=16 cells for S161A. Cells analyzed came from three independent experiments (one-way ANOVA with Tukey post hoc multiple comparisons test). (E-F) Quantification of PM association for transfected GFP-PLEKHA5^FL^ (E) and PLEKHA5^WW-H-BP-PH^-GFP (F) using PM-targeted proximity biotinylation. Representative western blots are shown at top with quantification of GFP signals in the streptavidin pulldowns at bottom (n=4 biological replicates, one-way ANOVA with Tukey post hoc multiple comparisons test).

**Extended Data Figure 6. (related to Figure 3).**
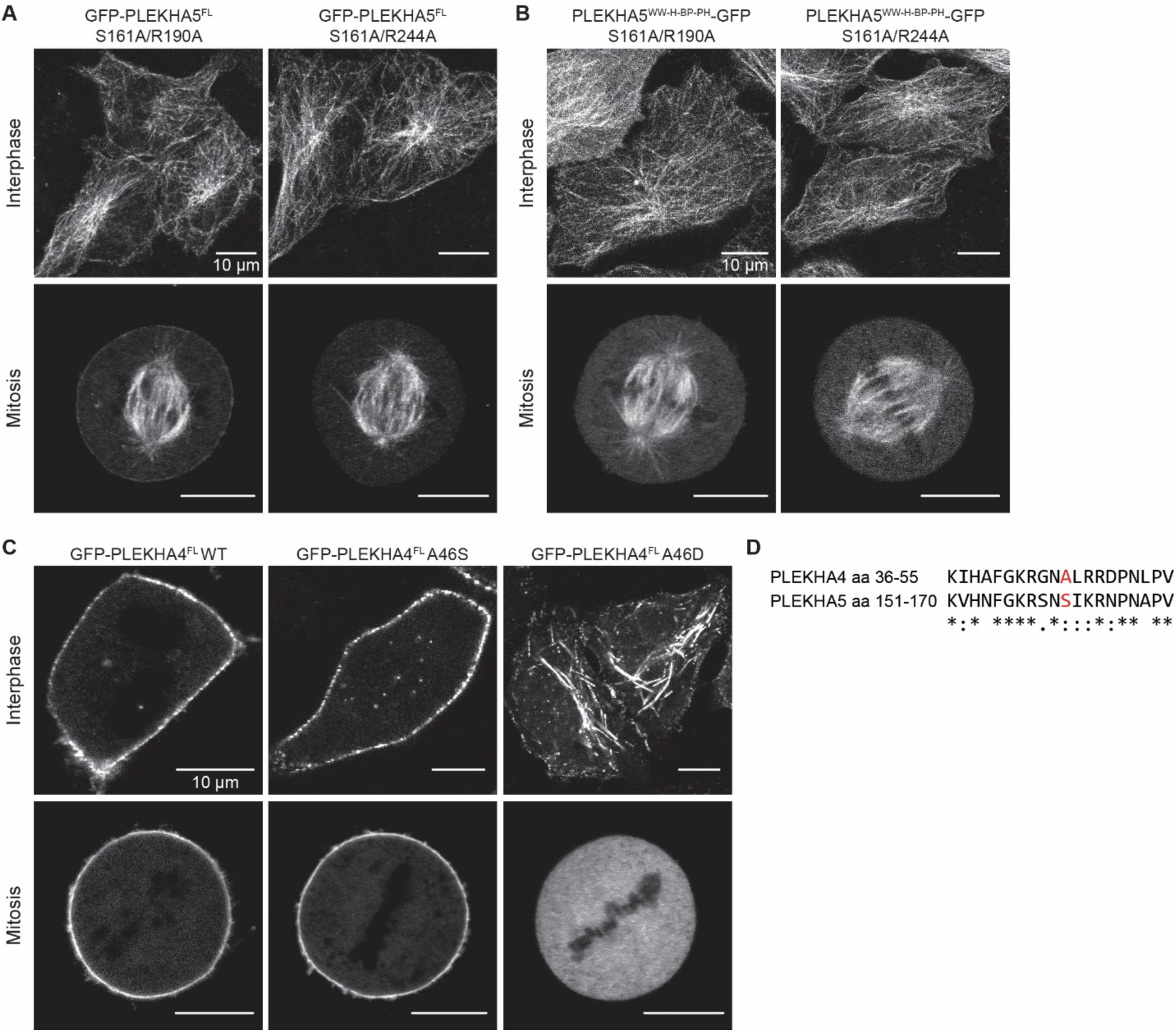
Requirement for PI(4,5)P_2_ binding in S161-mediated PM localization in additional PLEKHA5 constructs and conservation of S161-mediated PM localization in PLEKHA4. (A-B) Live-cell confocal microscopy of HeLa cells transiently expressing combinational mutants of S161A with either R190A or R244A in PLEKHA5^FL^-GFP (A) and PLEKHA5^H-BP-PH^-GFP (B). (C) Confocal microscopy of HeLa cells transfected with either the WT, A46S, or A46D forms of full-length PLEKHA4. (D) Clustal Omega alignment of amino acid sequences around A46 in PLEKHA4 and S161 in PLEKHA5.

**Extended Data Figure 7. (related to Figure 4).**
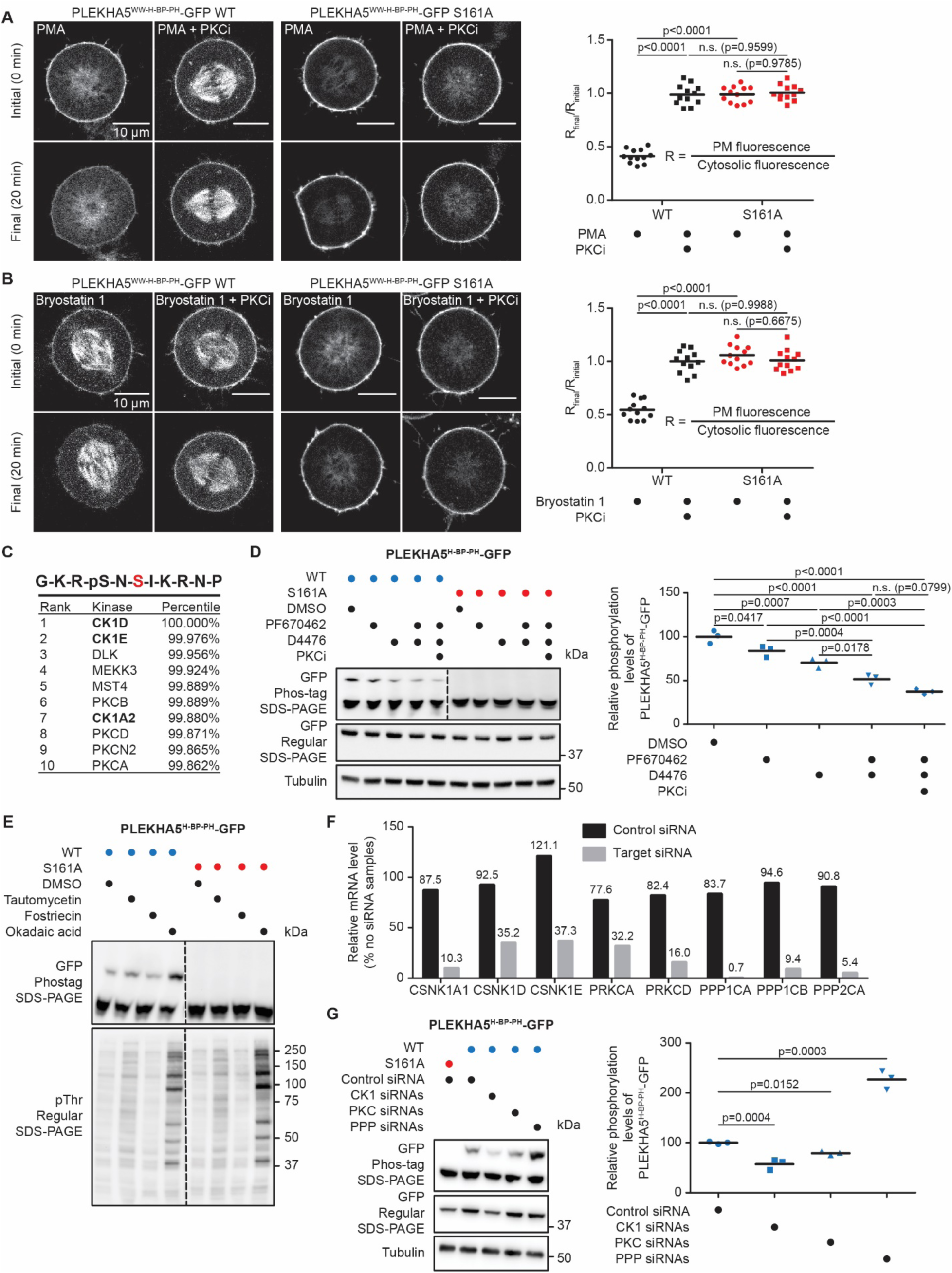
PKC and CK1 kinase activities contribute to S161 phosphorylation. (A-B) PLEKHA5^WW-H-BP-PH^-GFP displayed reduced PM localization upon activation of PKC by 100 nM PMA (A) or 200 nM Bryostatin 1 (B). Representative micrographs before and after agonist treatment for each condition are shown at left, with quantifications of the ratios of PM to cytosolic fluorescence (excluding microtubule signal) before and after treatment plotted on the right (n=12 cells from three independent experiments, one-way ANOVA with Sidak post hoc multiple comparisons test). (C) Top 10 kinase candidates for phosphorylating S161 (highlighted in red) if S159 serves as the priming phosphorylation site. (D) Inhibition of CK1 activity by 5 μM PF670462 or 100 μM D4476 decreased S161 phosphorylation. Shown are representative western blots and quantification for Phos-tag and regular SDS-PAGE analysis of PLEKHA5^H-BP-PH^-GFP (n=3 biological replicates, one-way ANOVA with Tukey post hoc multiple comparisons test). (E) Phos-tag SDS-PAGE analysis of S161 phosphorylation upon treatment of inhibitors of the PPP family of protein Ser/Thr phosphatases (Tautomycetin, Fostriecin, and Okadaic acid, each at 1 μM), probing with GFP and phospho-Thr antibodies. (F) RT-qPCR analysis of knockdown efficiency for siRNAs targeting CK1 isoforms (CSNK1A1, CSNK1D, CSNK1E), PKC isoforms (PRKCA, PRKCD), and PPP catalytic subunits (PPP1CA, PPP1CB, PPP2CA). (G) Knockdown of CK1 (triple knockdown of CK1α, CK1δ, and CK1ε) and PKC (double knockdown of PKCα and PKC8) decreased S161 phosphorylation, whereas depletion of PP1 and PP2A (triple knockdown of two PP1catalytic subunits and one PP2A catalytic subunit) increased S161 phosphorylation. Shown are representative Western blots and quantification for Phos-tag and regular SDS-PAGE analysis of PLEKHA5^H-BP-PH^-GFP (n=3 biological replicates, unpaired, two-tailed Student’s t-test).

**Extended Data Figure 8. (related to Figures 5 and 6).**
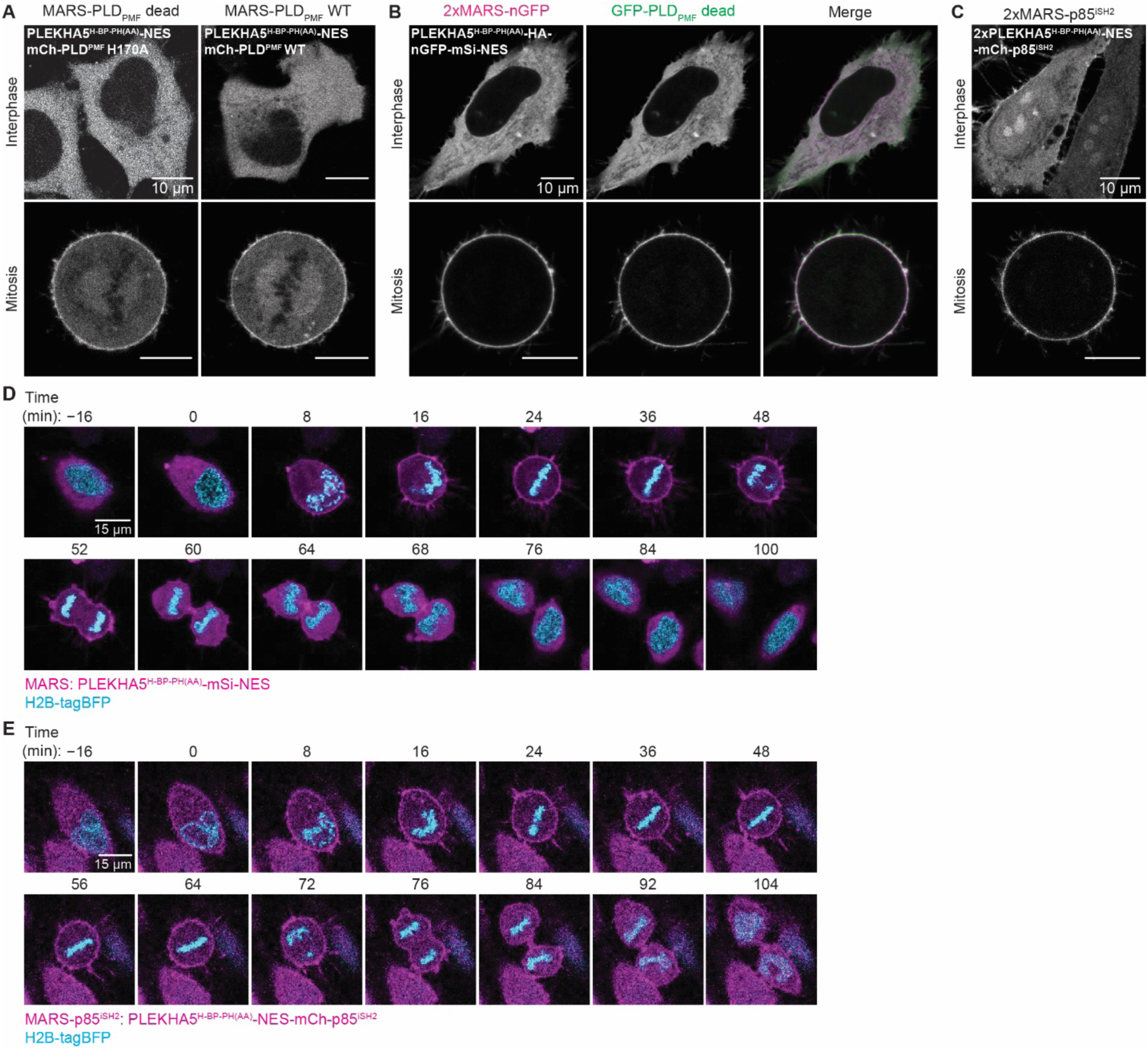
Evaluation of PM localization of additional MARS constructs for PLD and PI3K and time-lapse imaging of MARS and MARS-p85^iSH2^ during mitosis. (A) Live-cell confocal microscopy of HeLa cells transfected with the single MARS fusion to WT or catalytic dead (H170A) forms of PLD_PMF_ revealing only partial recruitment to the PM during mitosis. (B) Representative confocal micrographs of HeLa cells transfected with 2xMARS-nGFP (magenta) and H170A mutant of GFP-PLD_PMF_ (green) showing effective PM recruitment only during mitosis. (C) Live-cell confocal microscopy of HeLa cells transiently expressing the 2xMARS-p85^iSH2^ construct, revealing undesired partial PM localization during interphase. (D-E) Representative still images from time-lapse movies in asynchronous HeLa cells stably expressing H2B-tagBFP and transfected with either MARS (D) or MARS-p85^iSH2^ (E), and the PI(3,4,5)P_3_ marker GFP-Akt^PH^. Images display merged channels of H2B-tagBFP (cyan) and MARS constructs (magenta). Numbers indicate time before/after nuclear envelope breakdown (defined as t=0 min).

**Supplementary Video 1.** (related to Extended Data Figure 1B) Time-lapse confocal microscopy of asynchronous HeLa cells stably expressing H2B-mCherry and transiently transfected with PLEKHA5^WW-H-BP-PH^-GFP, during mitosis, with cells incubated at 37 °C and images acquired every 1 min. Left panel: H2B-mCherry; middle panel: PLEKHA5^WW-H-BP-PH^-GFP; right panel: merge (mCherry, magenta; GFP, green).

**Supplementary Video 2.** (related to Extended Data Figure 2C) Time-lapse confocal microscopy of asynchronous HeLa cells stably expressing H2B-tagBFP and transiently transfected with MARS, during mitosis, with cells incubated at 37 °C and images acquired every 2 min. Left panel: H2B-tagBFP; middle panel: MARS; right panel: merge (tagBFP, green; MARS, magenta).

**Supplementary Video 3.** (related to Extended Data Figure 2D) Time-lapse confocal microscopy of asynchronous HeLa cells stably expressing H2B-tagBFP and transiently transfected with 2xMARS-nGFP, during mitosis, with cells incubated at 37 °C and images acquired every 2 min. Left panel: H2B-tagBFP; middle panel: MARS; right panel: merge (tagBFP, green; 2xMARS-nGFP, magenta).

**Supplementary Video 4.** (related to Extended Data Figure 3E) Time-lapse video acquired by IncuCyte Live-Cell Analysis System of wild-type (WT) HeLa cells incubated at 37 °C with 5% CO_2_. Images were acquired every 15 min.

**Supplementary Video 5.** (related to Extended Data Figure 3E) Time-lapse video acquired by IncuCyte Live-Cell Analysis System of HeLa cells stably expressing MARS, with cells incubated at 37 °C with 5% CO_2_ and images acquired every 15 min.

**Supplementary Video 6.** (related to Extended Data Figure 3E) Time-lapse video acquired by IncuCyte Live-Cell Analysis System of HeLa cells stably expressing 2xMARS-nGFP, with cells incubated at 37 °C with 5% CO_2_ and images acquired every 15 min.

**Supplementary Video 7.** (related to Figure 6E and Extended Data Figure 8D) Time-lapse confocal microscopy of asynchronous HeLa cells stably expressing H2B-tagBFP and transiently transfected with MARS and the PI(3,4,5)P_3_ marker GFP-Akt^PH^, during mitosis, with cells incubated at 37 °C and images acquired every 4 min. The left video displays merged channels of H2B-tagBFP (cyan) and MARS (magenta), and the right video displays merged channels of H2B-tagBFP (cyan) and GFP-Akt^PH^ (yellow).

**Supplementary Video 8.** (related to Figure 6F and Extended Data Figure 8E) Time-lapse confocal microscopy of asynchronous HeLa cells stably expressing H2B-tagBFP and transiently transfected with MARS-p85^iSH2^ and the PI(3,4,5)P_3_ marker GFP-Akt^PH^, during mitosis, with cells incubated at 37 °C and images acquired every 4 min. The left video displays merged channels of H2B-tagBFP (cyan) and MARS-p85^iSH2^ (magenta), and the right video displays merged channels of H2B-tagBFP (cyan) and GFP-Akt^PH^ (yellow).

